# SKAP2 as a new regulator of oligodendroglial migration and myelin sheath formation

**DOI:** 10.1101/2021.04.22.439972

**Authors:** Julia Ghelman, Laureen Grewing, Farina Windener, Stefanie Albrecht, Alexander Zarbock, Tanja Kuhlmann

**Author notes:** Corresponding author: Tanja Kuhlmann, Institute of Neuropathology, University Hospital Muenster, Pottkamp 2, 48149 Muenster, Germany.

## Abstract

Oligodendroglial progenitor cells (OPC) are highly proliferative and migratory bipolar cells, which differentiate into complex myelin forming and axon ensheathing mature oligodendrocytes during myelination. Recent studies indicate that the oligodendroglial cell population is heterogeneous on transcriptional and functional level depending on the location in the CNS. Here, we compared intrinsic properties of OPC from spinal cord and brain on functional and transcriptional level. Spinal cord OPC demonstrated increased migration as well as differentiation capacity. Moreover, transcriptome analysis revealed differential expression of several genes between both OPC populations. In spinal cord OPC we confirmed upregulation of SKAP2, a cytoplasmatic adaptor protein known for its implication in cytoskeletal remodelling and migration in other cell types. Recent findings suggest that actin dynamics determine not only oligodendroglial migration, but also differentiation: Whereas actin polymerization is important for process extension, actin destabilization and depolymerization is required for myelin sheath formation. Downregulation or complete lack of SKAP2 in OPC resulted in reduced migration and impaired morphological maturation in oligodendrocytes. In contrast, overexpression of SKAP2 as well as constitutively active SKAP2 increased OPC migration suggesting that SKAP2 function is dependent on activation by phosphorylation. Furthermore, lack of SKAP2 enhanced the positive effect on OPC migration after integrin activation suggesting that SKAP2 acts as modulator of integrin dependent migration. In summary, we demonstrate the presence of intrinsic differences between spinal cord and brain OPC and identified SKAP2 as a new regulator of oligodendroglial migration and sheath formation.

**Significance statement:** OPC play an important role in many still incurable diseases, such as multiple sclerosis, leukodystrophies or neurodegenerative diseases. Their heterogeneity in different CNS regions has recently been identified. Here, we observed increased migration and differentiation capabilities of OPC isolated from the spinal cord compared to brain OPC and confirmed differences in the transcriptome between these two cell populations. Furthermore, we identified SKAP2 as potential modulator of actin dynamics in oligodendrocytes. Whereas knockdown or lack of SKAP2 impairs migration and myelin sheath formation in mouse and human oligodendrocytes, overexpression of wildtype or constitutively active SKAP2 enhances the migratory capacity of murine oligodendrocytes. In summary, we present SKAP2 as modulator of cytoskeletal dynamics regulating OPC migration, differentiation and myelin sheath formation.

## Introduction

Oligodendrocytes are the myelin forming cells of the CNS. During CNS development oligodendroglial progenitor cells (OPC) are generated in three waves at different anatomical sites in spinal cord and brain (Kessaris *et al*, 2006). OPC migrate through the brain along blood vessels (Tsai *et al*, 2016), start to extent multiple and branched processes and finally form the myelin sheath which is prerequisite for saltatory conduction of action potentials. Recently published studies demonstrated that oligodendrocytes (OL) are heterogeneous on transcriptional levels and that individual oligodendroglial subtypes dominate in certain CNS regions (Marques *et al*, 2016; van Bruggen *et al*, 2017); however it is unknown whether these differences in transcriptional levels also result in functional differences. Previously, several studies focusing on OPC derived from either white or grey matter found differences regarding proliferation (Hill *et al*, 2013; Lentferink *et al*, 2018), susceptibility to inflammatory cues (Lentferink *et al*, 2018) and differentiation capacity (Vigano *et al*, 2013). Moreover, other studies identified intrinsic functional and transcriptomic differences between spinal cord and brain-derived OPC (Bechler *et al*, 2015; Horiuchi *et al*, 2017; Marques *et al*, 2018).

The intracellular mechanisms driving oligodendroglial differentiation and myelin sheath formation have been extensively investigated during recent years (Franklin & Goldman, 2015; Simons & Nave, 2015; Kuhn *et al*, 2019), but only relatively little is known about the intracellular signaling cascades regulating oligodendroglial migration and how extracellular signals modify these intracellular pathways. Migration of OPC is controlled by extracellular molecules such as growth factors (e.g. FGFs, PDGF), guidance molecules (netrins, semaphorins) and chemokines (e.g. CXCL1), as well as by contact mediated mechanisms such as extracellular matrix molecules (for review see (de Castro *et al*, 2013)). A major factor for motility of OPC are the dynamics of the actin cytoskeleton (for review see (Thomason *et al*, 2020)). Recent studies indicate that actin polymerization is required for OPC migration and process extension whereas myelin sheath formation and axon wrapping is characterized by actin destabilization and depolymerization (Nawaz *et al*, 2015; Zuchero *et al*, 2015; Snaidero *et al*, 2017). However, regulators of intracellular signaling pathways orchestrating actin dynamics in both migrating and myelin sheath forming oligodendrocytes have not been described (for review see (Brown & Macklin, 2020)).

SKAP2 (*src-kinase associated protein 2*) is a cytoplasmic adapter protein that is expressed in several cell types such as lymphocytes, monocytes and neutrophils and which is required for global actin reorganization during cell migration (Alenghat *et al*, 2012; Boras *et al*, 2017). SKAP2 that has been first described in 1998 (Marie-Cardine *et al*, 1998), contains a pleckstrin homology (PH) domain, multiple tyrosine phosphorylation sites, C-terminal Src-Homology 3 domain (SH3 domain) and an N-terminal coiled-coil domain (Bureau *et al*, 2018). It interacts with different molecules implicated in integrin signaling events, including the adhesion and degranulation-promoting adaptor protein (ADAP) and RAP1-GTP–interacting adaptor molecule (RIAM) (Asazuma *et al*, 2000; Königsberger *et al*, 2010; Alenghat *et al*, 2012; Boras *et al*, 2017).

In this study, we observed increased migration and differentiation capabilities of OPC isolated from the spinal cord compared to brain OPC as well as differences in the transcriptome between these two cell populations confirming the presence of intrinsic differences between spinal cord and brain derived OPC. Furthermore, we demonstrate that knockdown or lack of SKAP2 impairs migration and myelin sheath formation in mouse and human oligodendrocytes whereas overexpression of wildtype or constitutively active SKAP2 enhances the migratory capacity of murine oligodendrocytes. SKAP2 is regulated by phosphorylation in OPC probably involving FYN (FYN Proto-Oncogene, Src Family Tyrosine Kinase) as direct upstream activator. Moreover, integrin activation via pRGD reverses the positive effect of SKAP2 on OPC migration. In summary, we identified SKAP2 as a new and important regulator of oligodendroglial migration and myelin sheath formation in mouse and human oligodendrocytes.

## Material and Methods

### Animals

For OPC isolation C57BL/6 mice or Skap-HOM mice (P6-9) were used. C57BL/6 were obtained from the Zentrale Tierexperimentelle Einrichtung (ZTE) of the university hospital of Münster. Skap-HOM mice (Togni *et al*, 2005) were kindly provided by Prof. Alexander Zarbock (Boras *et al*, 2017). All experiments were approved by the “Landesamt für Natur, Umwelt und Verbraucherschutz Nordrhein-Westfalen” (LANUV) and were performed according to the reference numbers 8.84.-02.05.20.12.286, 81- 02.05.50.17.017 and 84-02.04.2013.A029.

### Primary oligodendroglial cultures

Primary oligodendroglial progenitor cells (OPC) were isolated using the immunopanning method as described previously (Watkins *et al*, 2008). Briefly, dissociated mouse cerebrum or spinals cords were resuspended in panning buffer. To isolate OPC the single cell suspension was sequentially panned on two panning plates coated with anti-BSL 1 *Griffonia simplificonia* lectin (#L-1100, Vector Labs) and then incubated on a plate coated with anti-CD140a (#135902, Biolegend) for OPC selection. The adherent OPC were washed with PBS, scratched, and plated in 0.025% poly-L-lysine (PLL)-coated flasks (#P4707, Sigma). OPC-Sato medium was prepared by adding 100 U/mL penicillin, 10 µg/mL streptomycin (#P0781, Sigma), 292 µg/mL L-glutamine (#25030-024, Gibco), 100 µg/mL BSA (#A8806, Sigma-Aldrich), 100 µg/mL apo-transferrin (#T1147, Sigma-Aldrich), 60 ng/mL progesterone (#P8783, Sigma-Aldrich), 16 µg/mL putrescine (#P5780, Sigma-Aldrich), 40 ng/mL sodium selenite (#31966-021, Invitrogen), 60 µg/mL N-acetyl-cystein (#A9165, Sigma-Aldrich), 10 ng/mL biotin (#B4639, Sigma-Aldrich), 5 µg/mL insulin (#I1882, Sigma-Aldrich), 5 µM forskolin (#F6886, Sigma-Aldrich), Trace element B (#99-175, Cellgro) diluted in DMEM high glucose (#31966-021, Invitrogen) containing 2% B27 supplement with vitamin A (#12587010, Gibco). The cultures were maintained under proliferating conditions by addition of 10 ng/ml PDGF-AA (#100-13A, Pepro- Tech) and 5 ng/ml human NT3 (#450-03, Pepro-Tech). Differentiation of OPC was induced by replacing PDGF-AA by 10 ng/ml CNTF (#450-13, Pepro-Tech). The purity of oligodendroglial cultures was higher than 98 %.

### Lentiviral transduction of primary OPC

Production of murine ecotropic shRNA-containing lentiviruses was performed in HEK293T cell line (DSMB). HEK293T cells were transfected using Fugene HD transfection reagent in a 3:1 reagent : DNA ratio with packaging plasmid 2 µg psPAX2 (a gift from Didier Trono, Addgene, #12260), 1 µg murine ecotropic envelope plasmid pEnv(eco)-IRES-puro (Morita *et al*, 2000; Schambach *et al*, 2006) and 3 µg lentiviral transferplasmid. For knockdown experiments four pGFP-C-shLenti vectors containing different shRNAs against *Skap2* (Origene; see Table 1) were applied and scrambled shRNA was used as control (OriGene, #TR30021). For overexpression experiments, cDNA of SKAP2^WT^ and SKAP2^W336K^ (see Table 2) was transferred into pLEX_307 (Addgene #41392). For analysis of subcellular localization, lentiviral expression plasmid pSKAP2-mGFP (Origene #MR205468L2) was used.

**Table 1.**
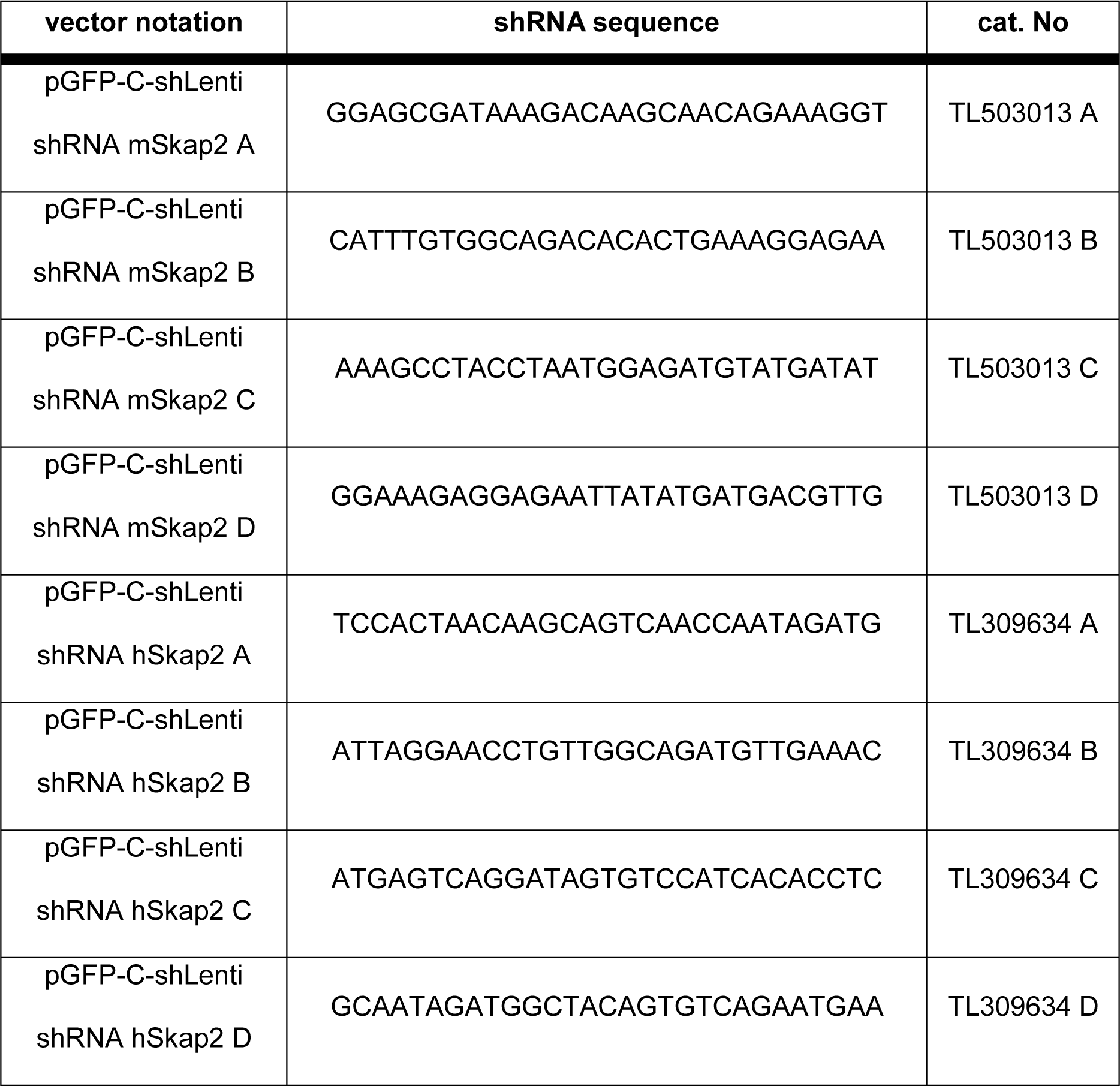
List of shRNA sequences used for knockdown of SKAP2 in murine OPC and hiOL.

**Table 2.** List of plasmids.

| no. | plasmid name | source | Purpose |
| --- | --- | --- | --- |
| 1 | psPAX2 | Addgene #12260 | Lentiviral packaging |
| 2 | pEnv(eco)-IRES-puro | Prof.Dr.Kitamura<sup>A</sup> | Lentiviral packaging |
| 3 | pLEX_307 (lentiviral backbone) | Addgene #41392 | Lentiviral transfer plasmid |
| 4 | pLEX_307_mCherry | Marc Ehrlich | Lentiviral expression plasmid with reporter |
| 5 | pGFP-C-shLenti | OriGene #TR30021 | Lentiviral knockdown, scrambled control |
| 6 | pGFP-C-shLenti (mSkap2) | OriGene #TL503013 A-D | Lentiviral knockdown of *Skap2* (mouse) |
| 7 | pGFP-C-shLenti (hSkap2) | OriGene #TL309634 A-D | Lentiviral knockdown of *SKAP2* (human) |
| 8 | pMaxGFP | Lonza | Control plasmid for transfection |
| 9 | pMIGII_Skap2-IRES-eGFP | Marc Boras (Boras et al., 2017) | cDNA template |
| 10 | pMIGII_Skap2<sup>W336K</sup>-IRES-eGFP | Marc Boras (Boras et al., 2017) | cDNA template |
| 11 | pLEX_SKAP2<sup>WT</sup> | This paper | Lentiviral transduction, SKAP2<sup>WT</sup> expression |
| 12 | pLEX_SKAP2<sup>W336K</sup> | This paper | Lentiviral transduction, SKAP2<sup>W336K</sup> expression |
| 13 | pSKAP2-mGFP | Origene #MR205468L2 | Lentiviral transduction, SKAP2 expression |
| 14 | pZFN-AAVS1_ELD | Addgene #159297 | Gene targeting |
| 15 | pZFN-AAVS1_KKR | Addgene #159298 | Gene targeting |
| 16 | pSON-rtTA | This paper | Gene targeting |

HEK293T supernatant containing virus particles was collected after 48 and 72 hours. Pooled supernatant was centrifuged (16-24 hours;12,300xg) using a centrifuge (5810R, Eppendorf) with an fixed angled-rotor (FA-45-6-30, Eppendorf) and the virus pellet was resuspended in OPC-Sato medium for 1 hour at 4°C on a shaker. Virus was either used freshly or was stored at -80°C for a maximum of 2 months. OPC were transduced with murine ecotropic shRNA-containing lentiviruses during replating. After 24 hours of incubation, medium was changed to remove viral particles. Transduction efficiency was tested 72 hours after viral transduction. For migration, cells were seeded in a density of 3000 cells/well in a 12-well plate and analyzed after 72 hours of 9 transduction. To analyze differentiation 80,000 cells/well were seeded in 6-well plates and medium was changed 24 hours after transduction to differentiation medium.

### Preparation of pRGD-coated plates

Solid pRGD (#F8141, Sigma) was diluted in H2O (1 mg/ml) and stored in the dark at room temperature. Shortly before coating, pRGD was diluted in a final concentration of 20 µg/ml in 20 mM HEPES in HBSS (#14175053, Gibco) containing 0.025 % poly-L-lysine (pRGD_HEP_). As control coating PLL was diluted in 20 mM HEPES in HBSS (PLL_HEP_). Plates were incubated for 2 hours at 37°C and washed with ddHM_2_O twice. Plates were always freshly prepared before use and not stored at 4° C.

### Preparation of matrigel-coated plates

Matrigel (#354263, Corning) was reconstituted by submerging the vial at 4°C over night and was diluted 1:5 in ice-cold knock-out DMEM (Cat.-No. 10829, Gibco) until material was evenly dispersed. The reconstituted matrigel was stored at -20°C upon use. Before plates were coated, matrigel was slowly thawed in knock-out DMEM 1:20 by pipetting up and down until matrigel was completely solved. Coated plates were stored at 4°C over night before use or for maximum 4 weeks.

### Stable integration of SON cassette in iPSC

For generation of an inducible SON- iPSC line, we adapted a one-in-all vector strategy for genomic integration to a save genomic harbour (Pawlowski *et al*, 2017). Human iPSCs were targeted with two zincfinger nucleases integrating the expression cassette containing the SON transcriptions factors (Ehrlich *et al*, 2017) under control of a Tet-On inducible promotor (TRE3G), a transactivator (rtTA) under control of CMV promoter as well as a puromycin resistance for selection into the adeno-associated-virus-site 1 (AAVS1) locus on intron 1 of chromosome 19 (Smith *et al*, 2008). pZFN-AAVS1_ELD, pZFN-AAVS1_KKR (both were a kind gift from Kosuke Yusa) and pSON-rtTA were transfected in a 1:1:1 ratio using FugeneHD. At day 2, selection with puromycin was initiated and surviving single cells were expanded to colonies and genotyped by locus and puromycin PCR to determine correct integration of the construct and homozygosity (Primer for genotyping, see Table 5). After induction with 1 µg/ml doxycycline (#D9891, Sigma) expression of *SOX10, OLIG2 and NKX2.2* was determined by RT-*q*PCR.

### Generation of smNPC from iPSC

Gene-targeted iPSC were differentiated into smNPC (referred to as NPC) applying a small molecule-based differentiation protocol (Reinhardt, Glatza, *et al*. 2013). Briefly, iPSC were cultured on mitomycin C (Sigma) inactivated mouse embryonic fibroblasts (EmbryoMax® PMEF, strain CF-1) in hESC medium consisting of Knockout DMEM (#10829-018, Invitrogen) with 20% Knockout Serum Replacement (#10828-028, Invitrogen), 1 mM β-mercaptoethanol (#31350-010, Invitrogen), 1% nonessential amino acids (#M7145, Sigma), 1% penicillin/streptomycin and 5 ng/ml bFGF. iPSC colonies were detached from MEF 3-4 days after splitting using 2 mg/ml collagenase IV. Colony pieces were collected by sedimentation and subsequently resuspended in hESC medium without bFGF but supplemented with 10 μM SB-431542, 1 μM dorsomorphin, 3 μM CHIR99021 and 0.5 μM purmorphamine and cultured as embryoid bodies (EBs) in petri dishes. Medium was replaced at day 2 with N2B27 medium consisting of 1:1 pre-mixed neurobasal medium (#21103-049) and DMEM/F12 (#21331-020, Gibco), 1% B27 supplement without vitamin A (#12587-010, Gibco), 0.5% N2 supplement (#17502-048, Gibco), 1% Pen/Strep (#P4333, Sigma), 2 mM L-glutamine (#G7513, Sigma) containing the same small molecules as before. On day 4, SB-431542 and dorsomorphin were removed from N2B27 medium and 150 μM ascorbic acid were added. On day 6, EBs were triturated into smaller pieces and plated onto matrigel-coated (#354263, Corning) 12-well plates in N2B27 medium supplemented with 0.5 μM purmorphamine, 3 μM CHIR99021 and 150 μM ascorbic acid.

### Human NPC culture

NPC were cultivated as described before (Reinhardt *et al*, 2013). In short, different NPC lines were stored in liquid nitrogen and cultivated in NPC medium on matrigel-coated wells. For passaging of NPC, cells were detached by adding accutase (#A6964, Sigma) at 37°C for 10 min. After detaching cell suspension was diluted 1:10 in split medium containing 1% BSA fraction V (#15260-037, Gibco) in DMEM low glucose (#D5546, Gibco) and cells were pelleted at RT at 200 rcf for 5 min. Cells were resuspended in pre-warmed N2B27 medium containing 1 µM SAG (#11914, Cayman Chemical Company), 3 µM CHIR99021 (#1386, Axon MedChem) and 150 µM L-ascorbic acid (#A4544, Sigma). Every second day medium was refreshed to ensure proper nutrient supply.

### Differentiation of hiOL

NPC lines with a stable integration of the inducible SON expression cassette were used for differentiation of hiOL. Generally, the protocol published by Ehrlich *et al*.(Ehrlich *et al*, 2017) was used and only small adjustments for the stable cell lines were made. In short, NPC were seeded with a density of 120,000 cells per well in a 12-well plate on matrigel. After 24 hours NPC were treated with N2B27 medium containing 1 µM SAG, 3 µM CHIR99021, 150 µM L-ascorbic acid and 1 µg/ml doxycycline (#D9891, Sigma) to initiate differentiation (day -2). Medium was switched to glial induction medium (GIM) supplemented with doxycycline to support differentiation (day 0). GIM contains 1% B27 supplement without vitamin A (#12587-010, Gibco), 0.5% N2 supplement (#17502-048, Gibco), 1% Pen/Strep (#P4333, Sigma), 2 mM L-glutamine (#G7513, Sigma), 1 µM SAG (#11914, Cayman Chemical Company), 200 µM L-ascorbic acid (#A4544, Sigma), 10 ng/ml T3 (#T6397, Sigma), 10 ng/ml IGF-1 (#100-11, Pepro Tech), 10 ng/µl PDGF-AA (#100-13A, Pepro- Tech), 10 ng/µl human NT3 (#450-03, Pepro-Tech) and 1:1000 Trace element B (#25022CI, Corning) diluted in DMEM/F12 (#21331-020, Gibco). At day 2 of differentiation medium was switched to glial differentiation medium (GDM) until end of differentiation. GDM contains 1% B27 supplement without vitamin A (#12587-010, Gibco), 0.5% N2 supplement (#17502-048, Gibco), 1% Pen/Strep (#P4333, Sigma), 2 mM L-Glutamine (#G7513, Sigma), 200 µM L-ascorbic acid (#A4544, Sigma), 60 ng/ml T3 (#T6397, Sigma), 10 ng/ml IGF-1 (#100-11, Pepro Tech), 10 ng/µl human NT3 (#450-03, Pepro-Tech), 200 µM dbcAMP (#D0627, Sigma) and 1:1000 Trace element B (#25022CI, Corning) diluted in DMEM/F12 (#21331-020, Gibco). At day 7 cells were plated to mouse laminin (#L2020, Sigma) coated well-plate formats and further differentiated until day 16 to day 21 in GDM. Medium was changed every second day to assure nutrient supply.

### Lentiviral transduction of hiOL

Production of human lentivirus was performed as described for murine virus (see above); however supernatant was only collected once after 72 hours post-transfection and 1.5 hours of ultracentrifugation (200,000 × g) was performed to pellet the virus. N2B27 medium was added to virus pellet and incubated without suspending at 4°C over night. Virus pellet was resuspended and stored in aliquots at -80°C. hiOL were transduced at day 8 of differentiation using 2.5 µL/mL virus. After 72 hours cells were checked for lentiviral plasmid expression.

### FACS of human hiOL

Cells were detached with accutase (#A6964, Sigma) at 37°C for 10 min., resuspended in split medium (1% BSA fraction V in DMEM low glucose) and centrifuged for 5 min. at 200 rcf. Cell number was determined using a Neubauer chamber. 98 µL of FACS buffer (0.5 % BSA in PBS) and 2 µL anti-O4-PE antibody (#130-118-978, Miltenyi Biotech) were added per 10^6^ cells. After incubation at 4°C for 10 min. cells were washed in 1 ml FACS buffer and centrifuged under same condition 13 as before. Pellet was again resuspended in appropriate volume of FACS buffer. Until analysis cells were stored on ice and were singularized before FACS procedure using a 35 µm nylon mesh cell strainer. Cells were sorted and analyzed using FACSMelody™ cell sorter (BD Bioscience) and the FACSChorus Software (Version 1.3 ©Becton Dickinson 2016).

### Cell migration assays

Cell migration was analyzed using the xCELLigence system that measures changes in impedance. In the upper chamber of a CIM-Plate 16 (ACEA Biosciences), 50,000 cells/well were plated. To analyze the migratory behaviour of mouse oligodendrocytes CIM-plates were coated with PLL or PLL/pRGD as described above. To stimulate OPC migration, 30 ng/ml PDGF-AA was added in the lower chamber. Impedance was measured every 15 min. for 24 hours and migration was quantified as relative cell index according to manufacturer’s protocol (xCELLigence, RTCA DP Analyzer, RTCA software 1.2, ACEA Biosciences). For each condition 4-6 technical replicates were included and experiments were performed with minimum 3 biological replicates. For analysis of xCELLigence data, a linear mixed model was applied (Shek & Ma, 2011). Statistical evaluation was performed with SPSS Statistics 24 (IBM). Mean relative cell index of technical replicates was calculated as percentage for each condition. The formula below describes the generation of individual growth curves (IGC) for both conditions including all biological replicates:

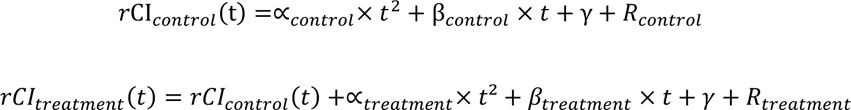

Based on our data those functions were computed for the control group (*r*CI_control_(t)) with γ describing the intercept, β the linear slope and α the square slope. R was the residual outcome of each individual replicate from the model. The lower R, the better did the curve describe the data set. For *r*CI_treatment_(t) the function was calculated in 14 dependency of *r*CI_control_(t). Cell index data were then transformed as curve function, which was statistically evaluated. Curves were compared to each other by computing a differential function and analyzing the parameters of initial slope (further referred as M*time) and slope over time (further referred as M*time2) and p values < 0.05 were considered significant.

For analysis of undirected migration, live cell imaging was performed (JuLI ™ Br, VWR); images were taken every 5 min. over 24 hours. Cells were manually tracked to assess velocity and migrated distance using ImageJ and the MtrackJ Plug-in (Meijering *et al*, 2012). For experiments using murine OPC 1000-2000 cells were seeded in a 12- well plate using either PLL or pRGD coating. For analysis of human OL 3000-4000 cells were seeded on wells coated with 5-10 µg/ml mouse-laminin (#L2020, Sigma) and 0.025 % poly-L-lysine (#P4707, Sigma) at day 7 of differentiation. hiOL were transduced at day 8 and migration was analyzed at day 11 (72 hours post transduction).

### Morphological analysis

Brightfield images were taken during differentiation using a Leica DMI6000 B inverted microscope. Pictures were taken 6, 24, 30 and 48 hours after onset of differentiation. At least 150 cells per time point and condition were classified by the following criteria: 0-2 processes (OPC), multiple branching (immature OL) or sheath formation (mature OL). Exemplary images for each condition are depicted in Fig. 3A. For evaluation of OPC process length under proliferating conditions, brightfield images were analyzed by manually measuring the longest protrusion per OPC using ImageJ.

### BrdU incorporation assay

Cells were cultured in a density of 3000 cells per coverslip and incubated with 20 µM BrdU (#B23151, Molecular Probes) in OPC-Sato medium for 6 hours. BrdU pulses were analyzed 0-6 hours, 24-30 hours, and 48-54 hours after 15 plating. Cells were fixed for 10 min. with 4% paraformaldehyde and permeabilized using 0.5% Triton-X for 10 min. After washing, cells were treated with cold 1M HCl for 10 min. and neutralized with Borat buffer for 10 min. Subsequent immunocytochemistry was performed as described below. For quantification 3 coverslips per time point were analyzed and percentage of BrdU^+^ cells was determined.

### 3D cell culture using nanofibers

Nanofiber inserts (Nanofiber solutions, 700nm, aligned; 24-Well Inserts, #242402) were coated using 0.01% w/v Poly-L-lysine-FITC labeled (#P3069, Sigma Aldrich) diluted in ddH_2_O at 4°C over night. Inserts were washed twice with PBS and cells were seeded in densities of 10,000 cells per well and cultured under proliferating conditions for 24 hours. Differentiation was initialized by addition of CNTF and cells were differentiated for 72 hours. Nanofibers inserts were fixated and stained similar to glass coverslips as described below (see Immunocytochemistry). Experiments were performed with in technical duplicates with 4 biological replicates (n = 4). For each replicate, minimum of 10 oligodendrocytes were imaged by using a confocal microscope in a 400x magnification. For evaluation of cell morphology and fiber ensheathment images were analyzed using a machine-based learning algorithm (in collaboration with Yu Kang T. Xu (Xu *et al*, 2019)).

### Immunocytochemistry (ICC)

For ICC cells were cultivated on 10 mm glass coverslips and fixed for 10 min. with 4% paraformaldehyde. Cells were permeabilized using 0.5 % Triton-X for 10 min. and washed three times with PBS. Cells were blocked with 5 % FCS (#S0115, Biochrome) in PBS and incubated with primary antibodies over night at 4 °C (Table 3). The next day cells were washed three times and afterwards Alexa-Fluor-conjugated secondary antibody was applied for 1 hour at room temperature. Subsequently, cells were washed three times with PBS and then mounted in Roti^®^ Mount FluorCare Dapi mounting medium (#HP20.1, Dako). Cells were visualised on a Zeiss LSM700 confocal microscope using ZEN software. To quantify the results of ICC at least 100-150 cells were quantified as percentage of DAPI positive cells for each condition and every independent experiment.

**Table 3.** List of antibodies.

| **primary antibody** | **species** | **dilution** | **company** | **cat. Number** | **purpose** |
| --- | --- | --- | --- | --- | --- |
| Anti-Tag(CGY)FP | rabbit | 1:2500 | evrogen | AB121 | IF |
| CD140a (PDGFrα) | mouse | 1:300 | Santa Cruz | sc-338 | IF |
| CD140a APA5 | rat | 1:300 | Biolegend | 135902 | Primary OPC |
| FYN | mouse | 1:100 | ThermoSc. | MA1-19331 | IF |
| GFP | rabbit | 1:1000 | Abcam | Ab6556 | IF |
| G3G4 (BrdU) | mouse | 1:4 | DSHB | AB 2314035 | IF |
| Ki-67 | rabbit | 1:100 | Abcam | AB16667 | IF |
| MBP | rat | 1:200 | abcam | Ab7349 | IF |
| NOGO-A | rabbit | 1:100 | Millipore | AB5664P | IHC/IF |
| OLIG-2 | rabbit | 1:100 | IBL | 18953 | IHC/IF |
| O4 | mouse | 1:100 | Merck | MAB345 | IF |
| O4-APC | mouse | 1:10 | Miltenyi<br>Biotec | #130-118-978 | FACS |
| SKAP55-R (C-9) | mouse | 1:50 | Santa Cruz | sc-398285 | IF/IHC |
| SKAP2 | rabbit | 1:1000 | ProteinTech | 12926-1-AP | IF/IHC |
| tGFP | rabbit | 1:5000 | evrogen | AB513 | IF |
| **Secondary antibody** | **Species** | **Dilution** | **Company** | **Cat. number** | **Purpose** |
| Alexa Fluor 488 anti-rabbit | goat | 1:250 | Jackson | 111-545-144 | IF |
| Cy3 anti-mouse | goat | 1:250 | Jackson | 715-165-150 | IF |
| Cy3 anti-rat | goat | 1:250 | Jackson | 112-165-167 | IF |
| Cy3 anti-rabbit | goat | 1:250 | Jackson | 111-165-144 | IF |
| IgG anti-rat | goat | 1:333 | Dianova | 112-005-003 | OPC |

### RNA isolation and quantitative real-time PCR (*q*PCR)

Total RNA from murine cells was isolated using peqGOLD Total RNA Kit (#12-6634, PeqLab). For RNA isolation of microarray probes, the Rneasy Mini Kit (#74104, Qiagen) was used. mRNA was transcribed into cDNA by reverse transcription reaction using High Capacity cDNA Transcription Kit (#4368814, Applied Biosystems) and the cDNAs were diluted to a final concentration of 0.75 ng/µl. All *q*PCR experiments were performed using Power SYBR™ Green PCR Master Mix (#4309155, Applied Biosystems). For 96-well microtiter plates StepOne Plus real time cycler (Applied Biosystems) and for 384-well microtiter plates a 7900 HT real-time cycler (Abi) was used. A standard *q*PCR protocol with 2 min. at 95 °C and 40 cycles of 30 sec. at 95 °C, 30 sec. at 55 °C and 30 sec. at 72 °C was applied. The melting curve of each sample was measured to ensure specificity of the amplified products. All samples were processed as technical triplicates. Data were normalized using Rplp0 as an internal control and analyzed by the Pfaffl ΔΔCt method (Livak & Schmittgen, 2001). Primers used for *q*PCR are provided in Table 4. Relative expression levels were calculated using the 2^−ΔΔCt^ method and normalized to biological reference samples.

**Table 4.**
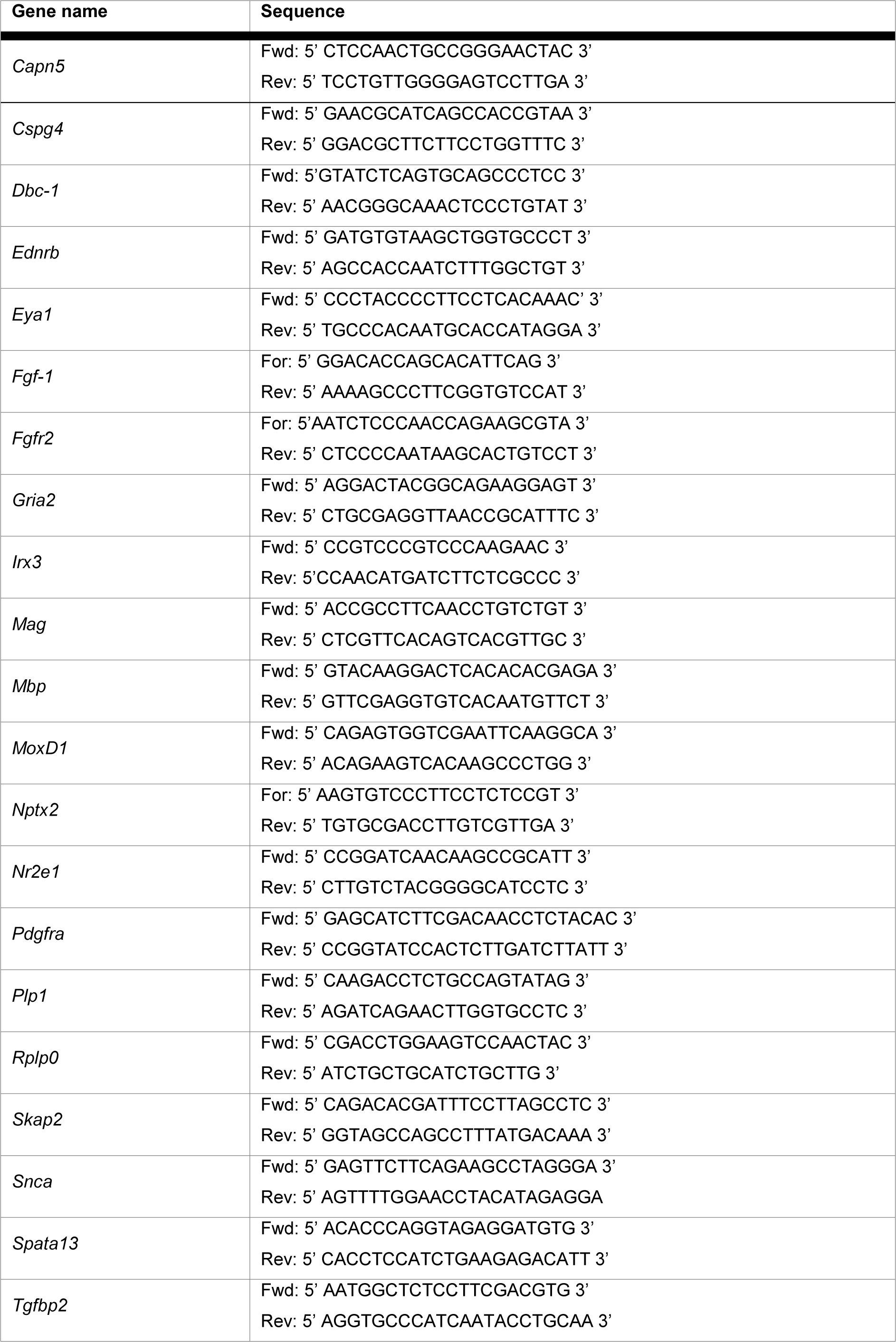
List of primers used for qPCR.

**Table 5.**
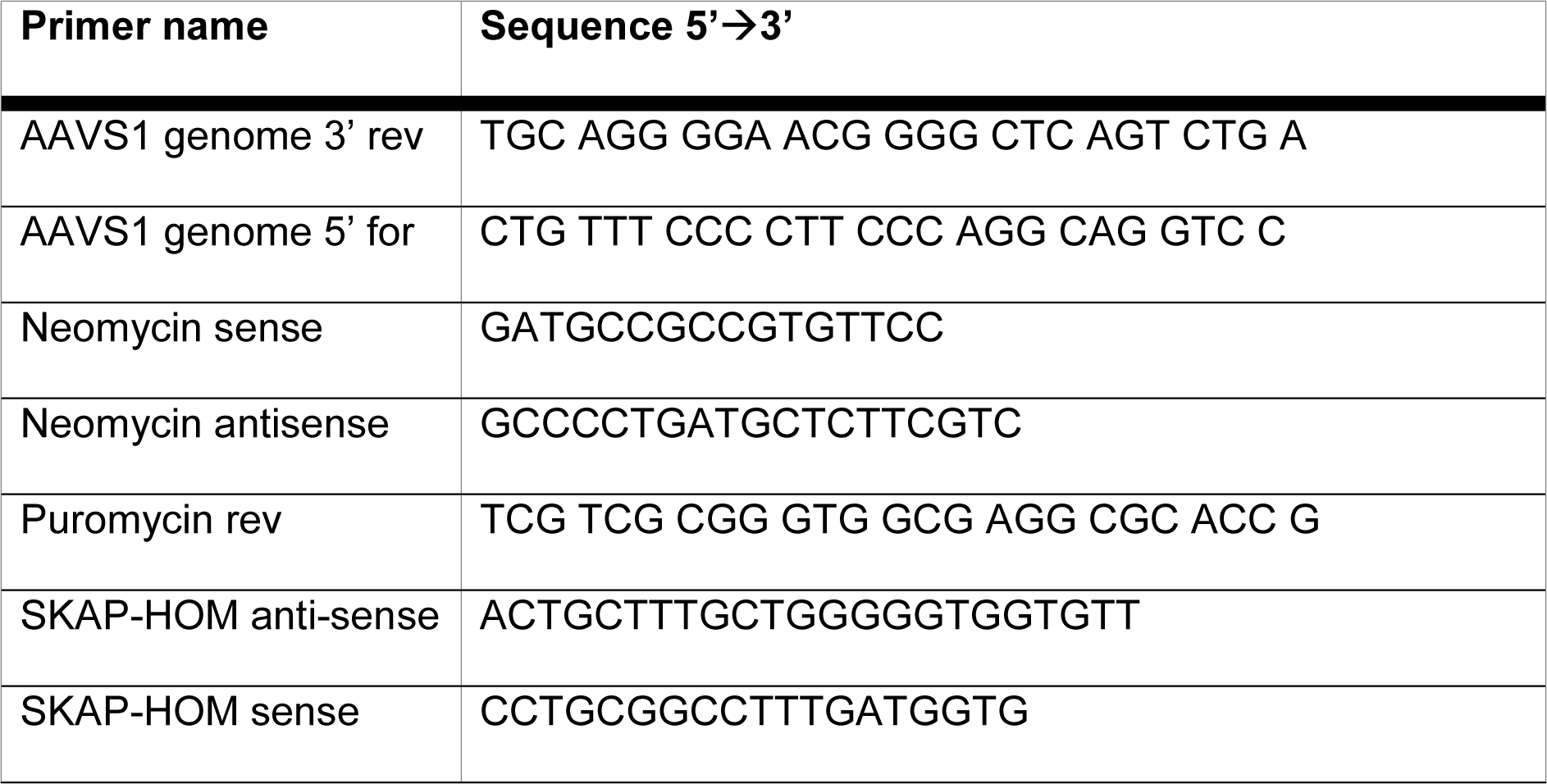
List of genotyping primers.

### Microarray

Total mRNA samples were used on MouseWG-6 v2.0 Expression BeadChips by Illumina. Differential expression was analyzed using direct hybridization analysis with quantile normalization, no background subtraction and the Illumina custom error model. cOPC served as reference group. Samples were collected after one passage and 48 hours of cultivation under proilferative conditions. Microarray was performed at the Core Facility Genomics at the University Hospital Muenster. Data analysis was performed using GenomeStudio (Illumina Inc. Version 2011.1).

### Experimental design and statistical analysis

All primary cell culture experiments were performed in technical triplicates and replicated minimum three times with biological independent samples (experiments performed with higher n-numbers are depicted in figure legends). Images used for morphological analysis or quantification of immunocytochemistry were taken randomly and from technical replicates (different wells or coverslips) in each independent experiment. Evaluation of images and cell counting were performed in a blinded fashion. All statistical analyses were performed using GraphPad Prism 5.03 (GraphPad Software Inc., San Diego, CA) or SPSS Statistics 24 (IBM). In text and figures the results are provided as means ± SEM if not mentioned otherwise. Experiments with n-numbers >7 were tested for Gauss distribution using the Anderson-Darling Test. According to this test, single comparisons were analyzed using two-tailed Student’*s t*-test. Multiple comparisons in the same data set were analyzed by Bonferroni-corrected (for selected groups) one- or two-way ANOVA tests. P values < 0.05 were considered significant (* p < 0.05, ** p < 0.01, *** p < 0.001).

## Results

### Higher migration and differentiation capacity of scOPC compared to cOPC

scOPC isolated from P6-8 mice displayed significantly increased undirected and directed migration capacities compared to cOPC as determined by live cell imaging and impedance measuring (Fig. 1A to E). Moreover, scOPC were characterized by longer protrusions compared to cOPC (Extendend data Fig. 1-1C). In contrast, we observed no differences in proliferation between the two cell populations as assessed by Ki67^+^ or BrdU-labelling (Extended data Fig. 1-1A-B and D). Oligodendroglial differentiation was induced by withdrawal of PDGF and addition of CNTF. After 48 hours we observed a significantly higher number of MBP^+^ mature scOPC than MBP^+^ cOPC (Fig. 1G to H). In line with this observation, the numbers of PDGFRα^+^ OPC were significantly reduced in scOPC compared to cOPC cultures (Fig. 1I). Furthermore, the number of MBP^-^ and PDGFRα^-^ immature oligodendrocytes was slightly, but significantly lower in scOPC compared to cOPC suggesting that not only the transition from PDGFRα^+^ OPC to immature oligodendrocytes but also from immature to MBP^+^ mature oligodendrocytes was accelerated in scOPC. Interestingly, no differences in the expression of *Mbp, Plp* or *Mag* were detected (Extenden data Fig. 1-1E to G). In summary, our results demonstrate that scOPC have a higher migration and differentiation capacity than cOPC.

**Fig. 1.**
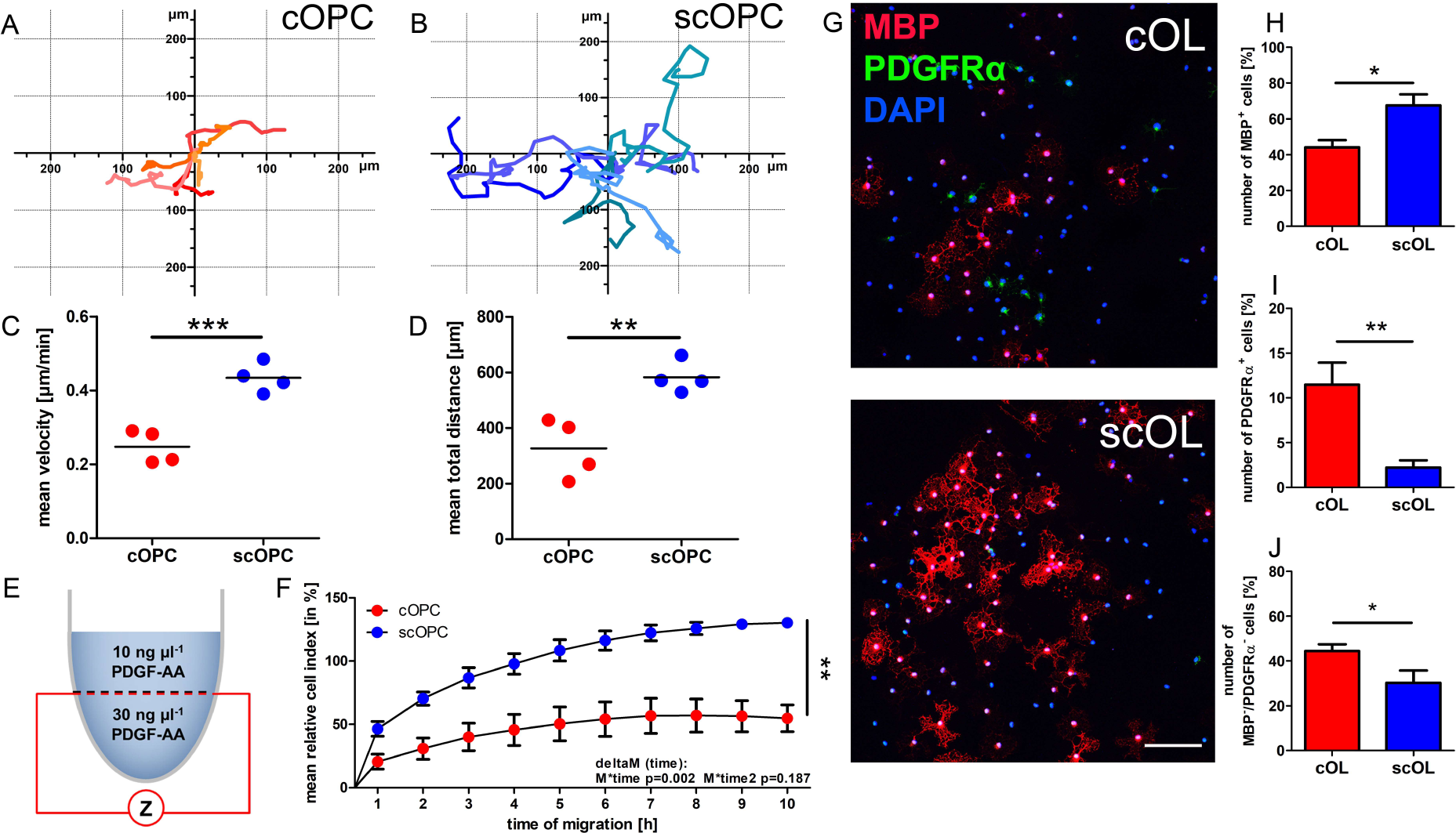
OPC from spinal cord have different migration and differentiation capacity compared to OPC from cerebrum. PDGFRα^+^ OPC were isolated from cerebrum and spinal cord. Exemplary cell tracking of scOPC and cOPC (A and B). Mean velocity and total migrated distance was increased in scOPC compared to cOPC after 24 hours of monitoring; statistic testing by unpaired t-test, p = 0.0008 for velocity and p = 0.0053 for total distance (C and D). Scheme to describe the experimental set-up to analyze directed migration towards a PDGF-AA gradient (E). scOPC displayed an increased migration rate towards PDGF-AA compared to cOPC, statistic testing by linear mixed model (F). ICC for MBP after 48 hours of differentiation, scale bar 100 µm (G). Quantification of MBP^+^, PDGFRα^+^ and PDGFRα^-^/MBP^-^ oligodendroglial lineage cells, p-values: 0.0102 for MBP^+^, 0.0048 for PDGFRα^+^ and 0.0489 for PDGFRα^-^/MBP^-^ (H to J).

### SKAP2 is differentially expressed in scOPC and cOPC

To gain insight how migration of scOPC is regulated on transcriptional level we compared the transcriptional profiles of scOPC and cOPC after induction of migration using 10 or 30 ng/ml PDGF-AA. Well-known OPC and OL marker genes, such as *Olig1, Olig2, PDGFRα, Mbp, and Plp*, were equally expressed in cOPC and scOPC (Fig. 2B). In contrast, patterning genes indicating a caudal (e.g. *Hoxa3, Hoxa5, Hoxa7, Hoxa9, Hoxc9*) or rostral (e.g. *Otx1, Foxg1, Moxd1*) origin of the cells were differentially expressed as expected. We identified 78 additionally differentially expressed genes (Extended data Table 2-1), 50 of them were significantly up- and 28 significantly downregulated in scOPC compared to cOPC after triggering of oligodendroglial migration by PDGF-AA (Fig. 2A). We selected 15 candidate genes for further validation by *q*PCR analyzing the same cDNA samples used for the microarrays (Extendend data Fig. 2-2A and B) and could confirm 10 candidate genes (including *Skap2*). Using a second set of independent cDNA samples from scOPC (n = 3) and cOPC (n = 3) 8 candidate genes were confirmed once more. *Skap2, Eya1, Capn5*, and *Nr2e1* were differentially expressed in both validation approaches (Extended data Fig. 2-2C to J). In subsequent experiments we focused on *Skap2* that has been described as modulator of actin dynamics and migration in other cell types (Alenghat *et al*, 2012; Ayoub *et al*, 2013; Tanaka *et al*, 2016). *Skap2* expression increased significantly during oligodendroglial differentiation *in vitro* (Fig. 2C) as well as in the corpus callosum of C57BL/6 mice during developmental myelination (Fig. 2D).

**Fig. 2.**
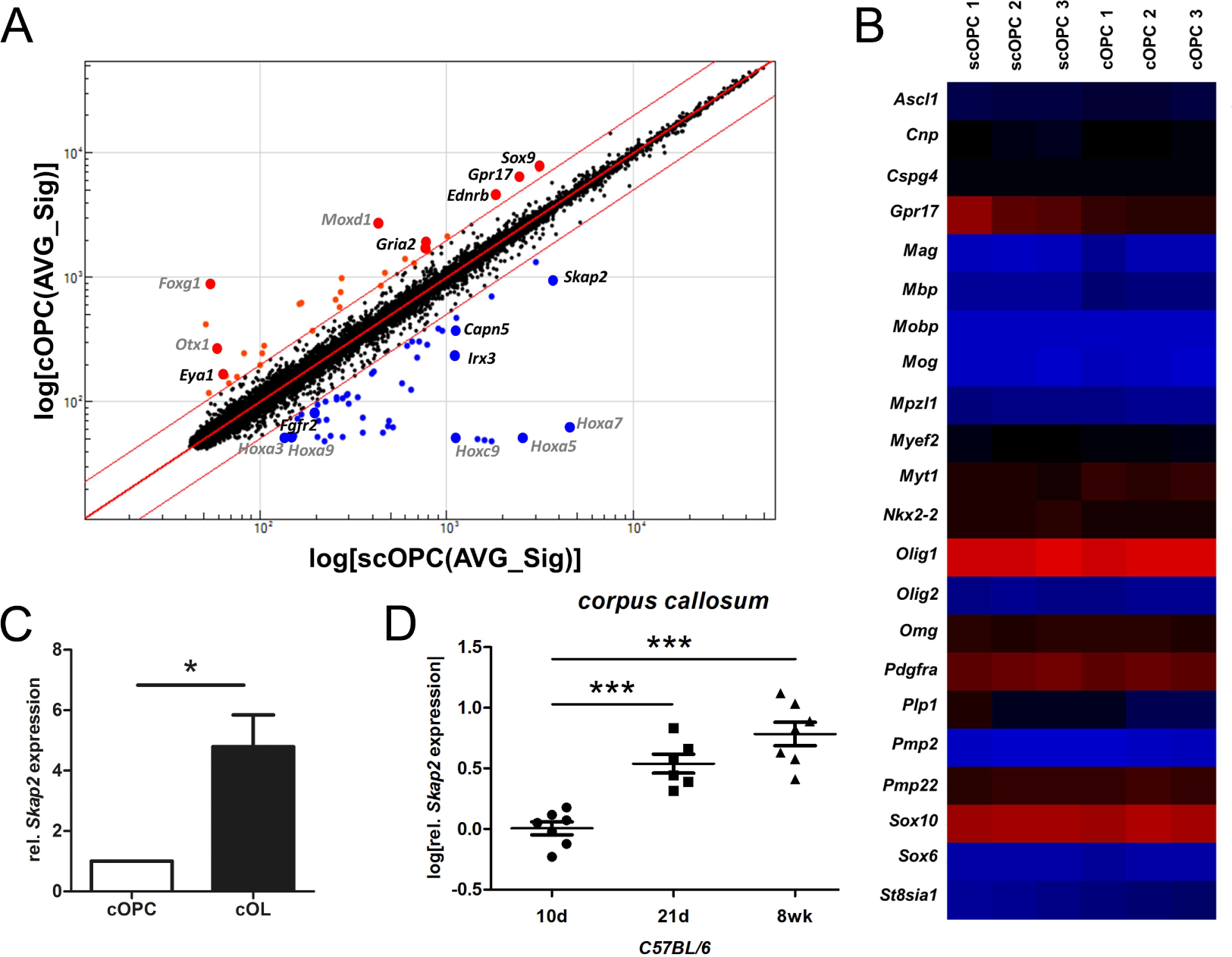
Differential gene expression between cOPC and scOPC and identification of *Skap2* upregulation during oligodendroglial differentiation *in vitro* and *in situ*. Dot plot of all analyzed genes depicts upregulation of patterning genes e.g. *Foxg1* for cOPC and *Hoxa7* for scOPC and further differential expressed genes between cOPC and scOPC. Red lines indicate threshold of 2-fold differential expression, red dots are genes upregulated in cOPC and blue dots upregulated in scOPC (A). Heat map showing selected oligodendroglial genes indicates no significant differences in expression between scOPC (n = 3) and cOPC (n = 3) (B). *Skap2* gene expression is upregulated during oligodendroglial differentiation *in vitro*, statistical testing by unpaired t-test with p = 0.007 (C) and myelination of the corpus callosum *in vivo*; 1-way ANOVA p < 0.0001, Bonferroni multiple comparison test (D).

### Knockdown or lack of SKAP2 reduces myelin sheath formation

To further examine whether SKAP2 has an impact on differentiation and myelin sheath formation we downregulated *Skap2* in oligodendrocytes using shRNAs and lentiviral transduction. Transduction efficiencies were in average 80 % resulting in a remaining *Skap2* expression of approximately 30 % in OL^Skap2sh^ compared to OL^scr^ after 48 hours of differentiation (Extended data Fig.3-1). We analyzed process and myelin sheath formation in differentiating oligodendrocytes using a morphological score (Fig. 3A). After knockdown of SKAP2 we observed reduced formation of myelin sheaths in cOL (Fig. 3B) and scOL (Extended data Fig. 3-1G), however expression of *Mbp* and *Plp1* were not affected (Extended data Fig. 3-1C to F). The use of a reporter plasmid in our knockdown experiments allowed us to analyze specifically transduced cells (Fig. 3C). The number of sheath forming tGFP^+^ cells was significantly reduced in cOL^Skap2sh^ compared to cOL^scr^ (Fig. 3D) whereas the number of cOL^Skap2sh^ with multiple branches was significantly increased compared to cOPC^scr^ (Fig. 3E). We did not observe a difference in the number of tGFP^+^O4^+^ or tGFP^+^MBP^+^ oligodendrocytes, further confirming that downregulation of SKAP2 does not affect the differentiation into O4^+^ and MBP^+^ mature oligodendrocytes (Extended data Fig. 3-2). We repeated the experiments by isolating oligodendrocytes from WT and SKAP2-deficient mice (Togni *et al*, 2005). cOL^SKAP2-/-^ displayed branched processes and a reduced number of myelin sheath forming cells compared to cOL^WT^ (Fig. 3F to G) similar to observations in scOL (Extended data Fig. 3-3F to G). Neither in scOL nor cOL lack of SKAP2 affected differentiation as determined by immunocytochemistry for MBP as well as *q*RT-PCR for *Mbp* and *Plp* respectively (Extended data Fig. 3-3A to D and Fig.3-4A to D). To analyze morphological differentiation in more detail in a 3D environment we cultivated OPC from wildtype and SKAP2^-/-^ on PLL-FITC coated nanofibers for 48 hours. scOPC, but not cOPC aligned and wrapped around nanofibers in a reproducible manner; therefore we performed our subsequent experiments with scOPC. Using an unbiased machine-based learning algorithm (Xu *et al*, 2019) we found an increased number of processes and total process length per cell in SKAP2^-/-^ oligodendrocytes supporting our previous findings from the 2D cultures (Fig. 3H to J). In summary, lack of SKAP2 prevented sheath formation and led to a rather branched morphology of differentiated oligodendrocytes compared to wildtype cells.

**Fig. 3.**
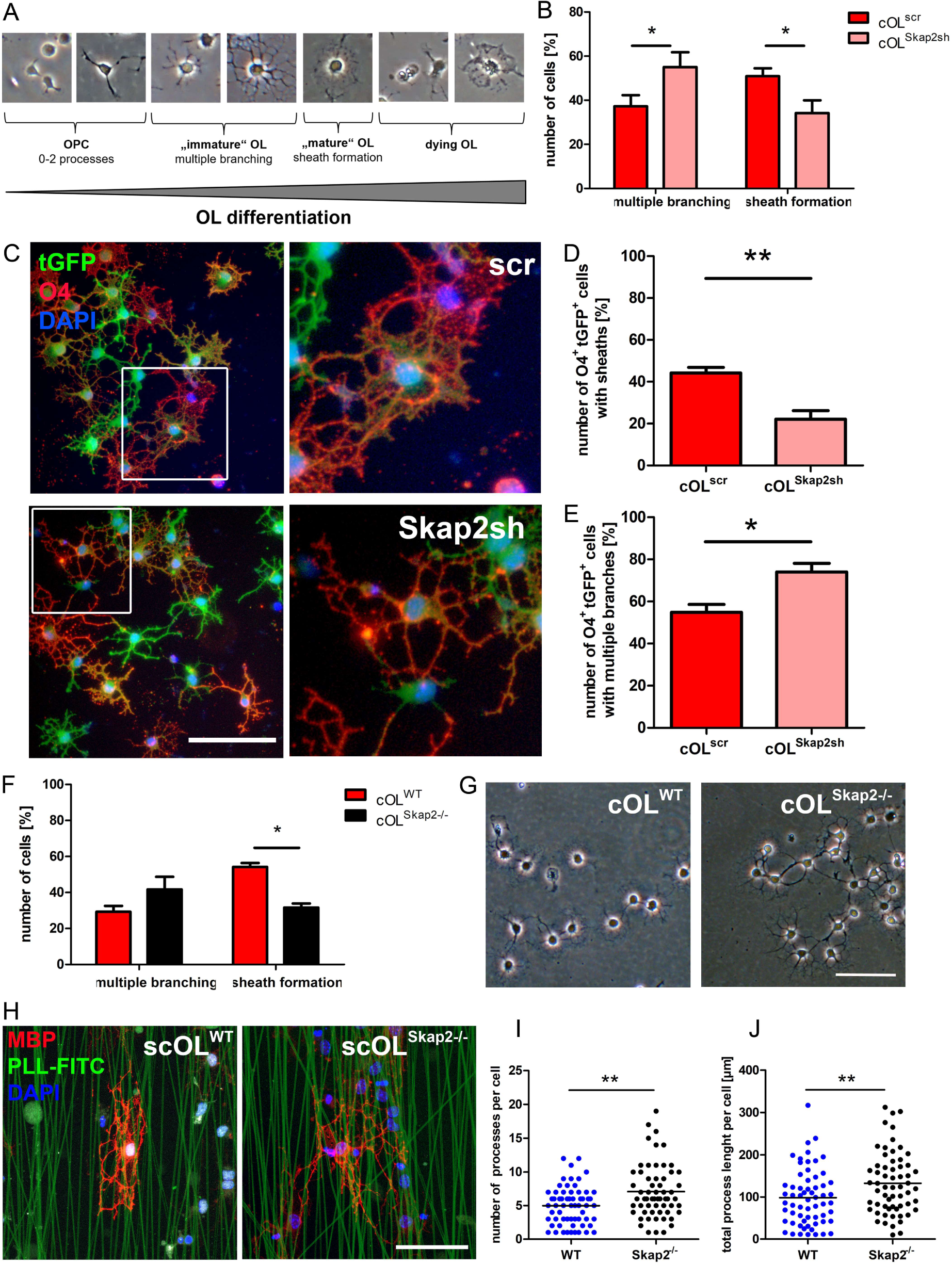
Knockdown and knockout of SKAP2 in oligodendrocytes impairs morphological maturation and myelin sheath formation. Typical morphology of differentiating oligodendrocytes is presented (A); for morphological analysis using brightfield images cells were categorized as multiple branching or sheath forming OL. cOPC transduced with *Skap2* shRNA displayed a reduced number of sheath forming cells (cOL: 50.89% ± 3.6 vs. 34.12% ± 5.8) and more cells remained as multiple branched cells (cOL: 37.23% ± 5.1 vs. 55.06% ± 6.7). Statistical analysis was performed with Two-way ANOVA and Bonferroni post-test with * p < 0.05 (n = 4) (B) Morphology of transduced tGFP^+^ and O4^+^ cOL was evaluated. Scale bar 100 µm (C). Downregulation of *SKAP2* results in a decreased number of sheath forming cells with 44.24% ± 2.7 in cOL^scr^ compared to 22.08% ± 4.2 in cOL^Skap2sh^ (D) and a higher number of cells with multiple branches with 54.87% ± 3.8 in cOL^scr^ and 74.01% ± 4.1 in cOL^Skap2sh^ (E). Quantifications demonstrate significantly reduced numbers of cOL^*Skap2*-/-^ forming sheaths compared to cOL^WT^ (31.63% ± 2.2 compared to 54.2% ± 2.1) (F). Brightfield images display reduced numbers of cOL^*Skap2*-/-^ forming sheaths after 48 hours, scale bar 100 µm (G). Exemplary image of scOL^WT^ and scOL^SKAP2-/-^ after 72 hours differentiation on PLL-FITC labeled 3D nanofibers. Scale bar 50 µm (H). Results of individual cell image analysis by machine-based learning algorithm are shown for number of processes per cell, unpaired t-test p = 0.0011 (I) and total process length per cell, unpaired t-test, p = 0.0079 (J).

### Knockdown or lack of SKAP2 reduces migration

To further elucidate the functional role of SKAP2 for migration we downregulated *Skap2* in migrating OPC (Extended data Fig. 4-1). Downregulation of Skap2 resulted in reduced undirected migration velocity and total distance in cOPC and scOPC as determined by live imaging (Fig. 4A to D). Due to limited cell numbers after transduction, no analysis of directed migration using the xCELLigence system was performed. Instead we examined the migratory behaviour of scOPC and cOPC derived from SKAP2^-/-^ mice. First, we analyzed the migration behavior of SKAP2-deficient cOPC (cOPC^SKAP2-/-^) and scOPC (scOPC^SKAP2-/-^) by cell tracking. Lack of SKAP2 in cOPC and scOPC resulted in slightly, but not significantly reduced undirected migration (Fig. 4E to H). Analysis of directed migration towards PDGF-AA using impedance measurements revealed decreased migration in SKAP2-deficient cOPC (Fig. 4I), but not scOPC (Fig. 4J). Lack of SKAP2 did not affect oligodendroglial proliferation in scOPC or cOPC as determined by Ki67 staining excluding decreased proliferation as cause for changes in impedance measurements (Extended data Fig. 3-3E and Fig. 3-4E). In summary, the combined data from knockdown and knockout experiments demonstrate that SKAP2 regulates oligodendroglial migration and suggest that constitutive lack of SKAP2 in OPC is at least partly compensated.

**Fig. 4.**
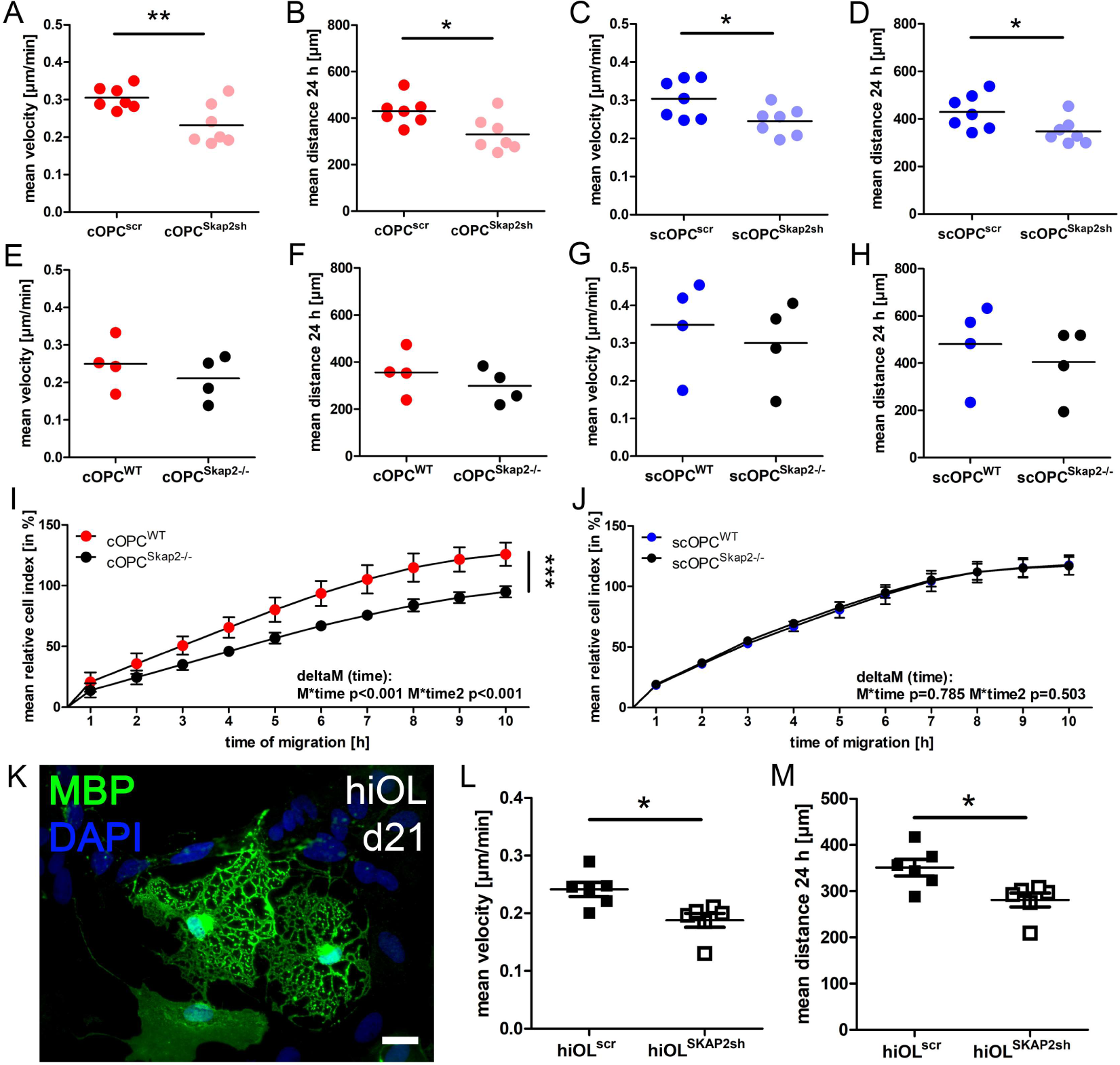
Lack of *SKAP2* decreased migration of murine OPC as well as iPSC derived human oligodendrocytes (hiOL) Lentiviral knockdown of Skap2 led to a reduced migratory capacity in cOPC as well as scOPC (n = 7) (A to D). Mean velocity in cOPC decreased from 0.30 ± 0.01 to 0.23 ± 0.02 µm min^-1^, p = 0.009 (A) and in scOPC from 0.30 ± 0.019 to 0.24 ± 0.014 µm min^-1^, p = 0.0293 (C). Total migrated distance after 24 hours was also decreased in cOPC from 430.6 ± 22.64 to 330.4 ± 28.19 µm, p = 0.0169 (B) and in scOPC from 430.0 ± 27.69 to 347.7 ± 20.45 µm, p = 0.0342 (D). Undirected migration of SKAP2- deficient OPC indicated a trend of decreased mean velocity (p = 0.0585) and migrated distance in cOPC (p = 0.1335) (n = 4) as well as in scOPC (E to H). Directed migration towards a PDGF-AA gradient was significantly decreased in cOPC^Skap2-/-^ compared to cOPC^WT^ (n = 3) (I), but not in scOPC (J). Exemplary image of MBP^+^ hiOL at day 21 of differentiation, scale bar 20 µm (K). Evaluation of directed migration showed a decreased velocity and total distance of *SKAP2* shRNA treated hiOL compared to scrambled shRNA hiOL, p = 0.0107, n = 6 (L and M).

### Knockdown of SKAP2 reduces migration of human iPSC derived oligodendrocytes *in vitro*

To determine whether SKAP2 is also relevant for the migration of human oligodendrocytes, we analyzed *SKAP2* in human induced pluripotent stem cell derived oligodendrocytes (hiOL) (Ehrlich *et al*, 2017). We confirmed an upregulation of SKAP2 in hiOL compared to undifferentiated iPSC derived NPC (Extended data Fig.4-2A). Moreover, migration of hiOL was analyzed by cell tracking after lentiviral knockdown using specific human *SKAP2* shRNAs (OriGene). 72 hours after infection we confirmed a knockdown efficiency of 37.9 % residual *SKAP2* expression (Extended data Fig.4- 2B). hiOL^SKAP2sh^ displayed a significant reduced velocity compared to hiOL^scr^ (Fig. 4K- M) demonstrating that downregulation of SKAP2 decreases migration not only in murine but also human oligodendrocytes.

### SKAP2 regulates migration via its phosphorylation

We further investigated whether increased SKAP2 expression or overexpression of an constitutive active form of SKAP2 influences oligodendroglial migration capacity as described for other cell types (Ayoub *et al*, 2013; Tanaka *et al*, 2016). We overexpressed wildtype SKAP2 (SKAP2^WT^) as well as a constitutive active mutant form of SKAP2 (SKAP2^W336K^) in murine cOPC (Fig. 5A). Overexpression of SKAP2^WT^ resulted in a significant increase in undirected migration; this was further enhanced when constitutive active SKAP2^W336K^ was overexpressed (Fig. 5B-D). FYN is also expressed in OPC (Sperber & McMorris, 2001) and it has been published that FYN is a direct upstream regulator of SKAP2 that phosphorylates SKAP2 at tyrosine 260 (Ayoub *et al*, 2013). To determine whether FYN colocalizes with SKAP2 we performed double immunofluorescent staining. We tested several SKAP2 antibodies (SKAP55-R (C-9), sc-398285 Santa Cruz; SKAP2 rabbit polyclonal AB, 12926-1-AP ProteinTech; SKAP2 rabbit polyclonal, HPA037468 Atlas Antibodies), however all of them showed a staining signal in SKAP2-deficient murine oligodendrocytes suggesting an unspecific signal. To circumvent this problem, we transduced wildtype OPC with a plasmid expressing monomeric GFP-tagged SKAP2 (pSKAP2mGFP). We detected SKAP2mGFP in the cytoplasm as well as in the leading edge of OPC as well as in the terminal processes of differentiating oligodendrocytes. Moreover, SKAP2mGFP distribution overlapped with F-actin indicating a close proximity of SKAP2 to the actin cytoskeleton (Fig. 5E). Co-staining with an antibody against FYN demonstrated colocalization of SKAP2 and FYN (Fig. 5F).

**Fig. 5.**
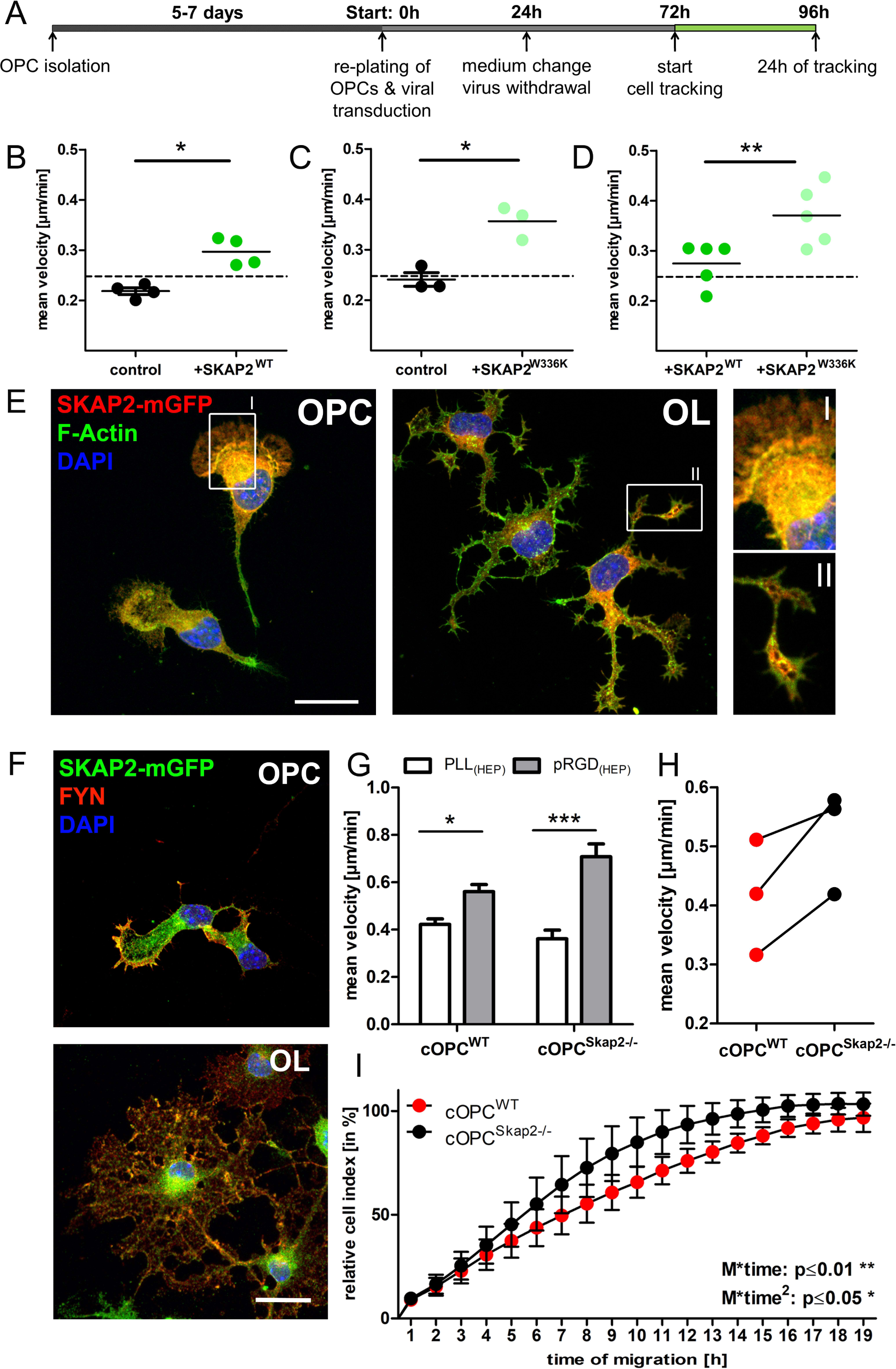
Function of SKAP2 in oligodendroglial migration is dependent on phosphorylation and SKAP2 colocalizes with FYN and F-Actin. Schematic overview of experimental set-up for evaluation of SKAP2 overexpression is presented (A). Overexpression of SKAP2^WT^ as well as SKAP2^W336K^ increased mean velocity of OPC compared to control, p-values 0.0138 and 0.0282 (B and C) and direct comparison between SKAP2^WT^ and SKAP2^W336K^ showed higher velocity of OPC expressing constitutively active SKAP2^W336K^, p-value 0.0024 (D). Expression of mGFP-tagged SKAP2 in OPC and differentiated oligodendrocytes showed an accumulation of SKAP2 in expanding processes and a colocalization with F-actin. Scale bar 20 µm (E). mGFP-tagged SKAP2 and FYN displayed a partial colocalization in processes and lammelipodia. Scale bar 20 µm (F). Mean velocity of OPC was increased on pRGD-coated surface compared to PLL in both cOPC^WT^ (p = 0.014) and cOPC^SKAP2-/-^ (p = 0.0062) evaluated by Student’s t-test (G). Undirected migration was increased on pRGD in cOPC^SKAP2-/-^ compared to cOPC^WT^ (H). Direct migration evaluated by xCELLigence was increased in cOPC^SKAP2-/-^ compared to cOPC^WT^ on pRGD coating evaluated by linear mixed model (I).

### SKAP2 is implicated in RGD-mediated oligodendroglial migration

In other cell types such as macrophages SKAP2 mediates integrin induced migration (Alenghat *et al*, 2012). Integrins are αβ-heterodimeric receptors located in the cell membrane and expressed by oligodendroglial cells in a stage-dependent manner (O’Meara *et al*, 2011). Integrins can either be activated by inside-out or outside-in mechanisms. Half of the integrins recognize and bind the peptide sequence Arg-Gly-Asp (RGD) leading to an outside-in integrin activation (Pierschbacher & Ruoslahti, 1984; Gailit & Ruoslahti, 1988; Nieberler *et al*, 2017). Cultivating cOPC^WT^ on a pRGD coated surface increased mean velocity of OPC significantly (Fig. 5G) suggesting a positive effect of integrin activation on oligodendroglial migration capacity. To determine whether SKAP2 is involved in RGD-mediated migration, we compared migration of WT and SKAP2-deficient cOPC on pRGD. Unexpectedly, cOPC from SKAP2-deficient mice showed an increased velocity and a stronger directed migration on pRGD coated wells compared to cOPC^WT^ (Fig. 5G-I). In summary, our data suggest that RGD-induced outside-in activation of integrins enhance migratory capacity of cOPC and that SKAP2 acts as modulator downstream of integrin induced migration.

## Discussion

In this study, we demonstrate that scOPC display increased migration and differentiation capacities but no differences in proliferation compared to cOPC. We confirmed differential gene expression between both OPC populations and identified SKAP2 as potential regulator of cytoskeletal dynamics in oligodendrocytes. Knockdown or lack of SKAP2 results in reduced oligodendroglial migration and myelin sheath formation, whereas overexpression and phosphorylation of SKAP2 increases oligodendroglial migration. Moreover, SKAP2 is associated with the actin cytoskeleton and implicated in pRGD-mediated migration of OPC.

OPC isolated from spinal cord and brain differ on a functional as well transcriptional level. scOPC and cOPC expressed typical caudal and rostral patterning genes, such as Hox genes in scOPC (Tumpel *et al*, 2009; Miguez *et al*, 2012; Philippidou & Dasen, 2013) and *Foxg1, Moxd1* and *Otx1* in cOPC (Schilling & Knight, 2001; Takuma & Carina, 2017; Tsuneoka *et al*, 2017) demonstrating that the developmental origin of the two cell types persisted *in vitro* comparable to descriptions in previous publications (Horiuchi *et al*, 2017; Marques *et al*, 2018). On functional level we observed an accelerated differentiation of scOPC into MBP^+^ cells compared to cOPC. This is in line with the observation by Bechler and colleagues who described longer internodes formed by rat spinal cord OPC compared to cerebrum OPC (Bechler *et al*, 2015). However, our results are in contrast to an earlier publication reporting lower proliferation and reduced MOG expression as well as myelin sheath formation in rat scOPC compared to forebrain (Horiuchi *et al*, 2017). These contradicting results might be at least partly explained by different species, different cell culture techniques or different assays to analyze proliferation, differentiation and myelin sheath formation. Interestingly, we did not observe a difference in *Mbp* mRNA expression between scOPC and cOPC despite significant differences in the number of MBP^+^ oligodendrocytes suggesting that accelerated differentiation of scOPC is not due to changes in myelin gene expression, but rather due to post-transcriptional modifications of myelin-associated genes or changes in cellular pathways regulating process and myelin sheath formation.

To further dissect the molecular pathways underlying the functional differences between scOPC and cOPC we analyzed RNA expression patterns of scOPC and cOPC. SKAP2 was among the differentially expressed genes and has been described as regulator of actin cytoskeletal dynamics in diverse immune cell types (Bourette *et al*, 2005; Zhou *et al*, 2011; Alenghat *et al*, 2012). Intrinsic *Skap2* expression was higher in scOPC compared to cOPC and associated with increased migratory capacity in scOPC. Whereas *Skap2* knockdown led to a similar reduction of migration in scOPC and cOPC, a constitutive lack of SKAP2 did only affect cOPC, but not scOPC suggesting that compensatory mechanisms may be more pronounced in scOPC. However, knockdown as well as knockout of *Skap2* prevented membrane spreading and myelin sheath formation in cOL and scOL in a comparable manner indicating that lack of SKAP2 cannot be compensated in the knockout situation.

Overall, the combined results from our knockdown, knockout and overexpression experiments demonstrate that SKAP2 is a positive regulator of migration in mouse and human oligodendrocytes similar to its function in immune cells and fibroblasts (Alenghat *et al*, 2012; Ayoub *et al*, 2013; Tanaka *et al*, 2016). In these cell types SKAP2 interacts with WASP (Tanaka *et al*, 2016) or SIRPα and ADAP (Alenghat *et al*, 2012) and mediates their recruitment to integrins regulating integrin-dependent cytoskeletal rearrangements and migration. In contrast, in glioblastoma cells SKAP2 acts as negative regulator of migration. Here, SKAP2 directly interacts with WAVE2 and cortactin, thereby preventing their association with ARP2/3, inhibiting actin assembly and subsequent migration (Shimamura *et al*, 2013) suggesting that the outcome of SKAP2 for migration depends on its binding partners and involved signaling cascades.

SKAP2 is an adaptor protein with many binding sites and a variety of binding partners regulating its activity and conformational changes (Zhou *et al*, 2011; Alenghat *et al*, 2012; Ayoub *et al*, 2013; Tanaka *et al*, 2016; Bureau *et al*, 2018). This is in line with our observation that SKAP2 acts generally as a positive regulator of oligodendroglial migration, but negatively affects OPC migration after integrin activation by stimulation with pRGD. Our results suggest that integrin-dependent and independent mechanisms regulate OPC migration and that both pathways are modulated by SKAP2. Future experiments focusing on protein interactions with respect to SKAP2 activation using PLA or Pull-down assays may help to dissect further the SKAP2 signalling network in oligodendroglial cells.

One mechanism of SKAP2 activation is phosphorylation of tyrosine-260 which is essential for cell migration of mouse embryonic fibroblasts; phosphorylation of SKAP2 by FYN positively affects migration and dephosphorylation by PTP-PEST results in defective migration (Ayoub *et al*, 2013). In our study overexpression of SKAP2^WT^ and SKAP2^W336K^ increased migration of OPC; however, the effect of SKAP2^W336K^ was significantly stronger than the effect of SKAP2^WT^ corroborating that phosphorylation of SKAP2 further promotes OPC migration. So far, it is unclear whether in oligodendrocytes regulation of SKAP2 activity is mediated by FYN and/or PTP-PEST as well. We confirmed that FYN colocalizes with SKAP2 in OPC supporting the hypothesis that SKAP2 and FYN interaction contributes to regulation of oligodendroglial migration. FYN has been identified to be involved in process outgrowth and ATP-induced migration of OPC (Klein *et al*, 2002; Bauer *et al*, 2009; Feng *et al*, 2015) as well as PDGF-AA dependent directed migration via cyclin-dependent kinase 5 (Cdk5) and WAVE2 (Miyamoto *et al*, 2008). Therefore, it can be hypothesized that in oligodendrocytes FYN and SKAP2 may be members of a common signalling network involving WAVE2 and directly targeting the actin cytoskeleton. The close proximity of SKAP2 and FYN to F-actin suggests further their involvement in actin dynamics and cytoskeletal rearrangements, which are both important prerequisites for cell migration as well as myelin sheath formation.

In migrating OPC dynamic formation and stabilization of processes is in balance with retraction and process destabilization. At early differentiation pre-oligodendrocytes become multipolar by stabilizing formed processes (Brown & Macklin, 2020). Subsequently, disintegration of cell processes by actin destabilization enables membrane spreading and formation of a 2-dimensional sheath structure which are typical hallmarks of OL differentiation *in vitro* (Barateiro & Fernandes, 2014; Thomason *et al*, 2020).

The regulation of the balance between actin stabilization and destabilization in processes of oligodendroglial cells is not fully understood. Our data demonstrate that SKAP2 is involved in the regulation of membrane spreading and myelin sheath formation; lack of SKAP2 impairs the transition from multipolar OL to sheath forming OL without affecting other hallmarks of maturation like MBP expression indicating a positive effect of SKAP2 on actin destabilization and subsequent process disintegration.

In oligodendrocytes actin polymerization is important for process extension and migration whereas actin destabilization and depolymerization is required for myelin sheath formation (Nawaz *et al*, 2015; Zuchero *et al*, 2015). Our results substantiate that SKAP2 is involved in both cellular processes. Similarly, mTOR has been identified as upstream regulator of actin reorganization affecting both, process extension as well as myelin sheath formation (Musah *et al*, 2020). Interestingly, the mTOR regulated proteome includes also FYN (Tyler *et al*, 2011) providing a potential link to upstream activation of SKAP2. Moreover, a signaling network involving N-WASP as key regulator of ARP2/3-driven actin polymerization and myelination has been described indicating another possible mechanism involving SKAP2 as adaptor (Katanov *et al*, 2020). Taken together, we hypothesize that SKAP2 is most important during phases of highest actin turnover and functions as adaptor protein, which associates with either F-actin directly mediating its stability or different binding partners modulating actin polymerization and depolymerization, respectively (Fig. 6). To understand to which extent SKAP2 affects actin reorganization in OPC and at different stages of oligodendroglial differentiation further experiments e.g. actin polymerization assays are required.

**Fig. 6.**
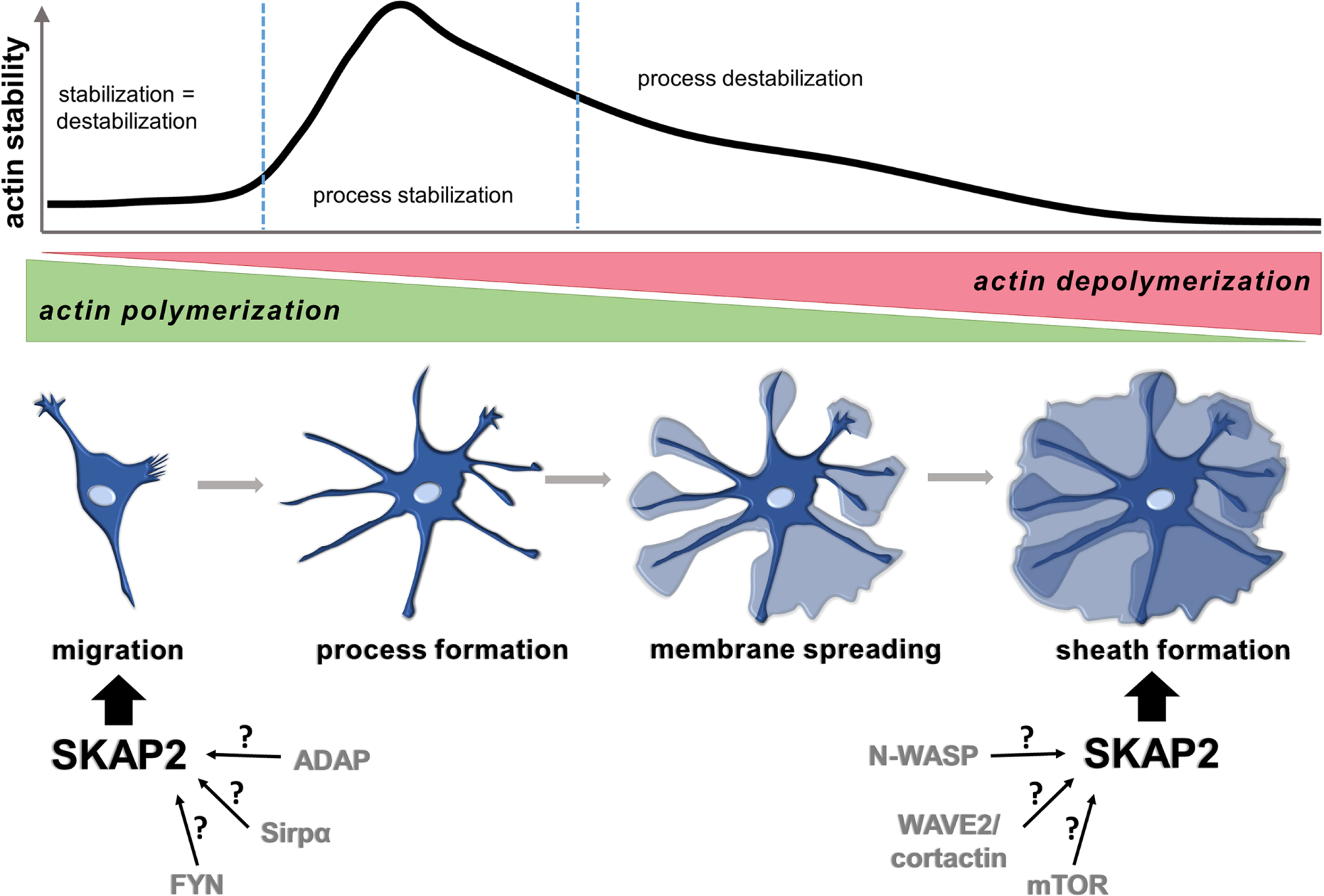
SKAP2 is involved in OPC migration as well as myelin sheath formation in oligodendrocytes. Schematic overview of actin dynamics in OPC and during oligodendroglial differentiation. Migrating OPC increased actin polymerization but low f-actin stability enables continuous formation and retraction of processes. With increasing actin stability formed processes can be sustained and differentiation is initiated. After multiple branches are formed actin stability decreases and increasing actin depolymerization facilitates membrane spreading and myelin sheath formation.

In conclusion, we demonstrated that OPC from brain or spinal cord have intrinsic properties with respect to migration and differentiation further supporting the notion of regional oligodendroglial heterogeneity. Furthermore, we identified SKAP2 as an important regulator of oligodendroglial migration and myelin sheath formation in mouse and human oligodendrocytes. To which extent SKAP2 contributes to OPC migration *in vivo* and under pathological conditions needs to be investigated in future studies. Further dissecting the molecular cascades regulating SKAP2 function might results in potential drug targets enhancing oligodendroglial migration and/or myelin sheath formation in demyelinating diseases such as multiple sclerosis.

## Acknowledgements

We thank Claudia Kemming and Elke Hoffmann for excellent technical support.

This work was supported by the Interdisciplinary Center for Clinical Research Münster (KuT3/012/15) and the German Research Foundation (Ku1477/11-1 and ZA 428/12- 1).

## Extended data legends

**Fig. 1-1** Proliferation and morphology of scOPC and cOPC

Double-staining with F-Actin and Ki67 and BrdU. Scale bars 100 µm (A). Quantification of Ki67^+^ nuclei did not reveal a difference in proliferative capacity of both OPC types; approximately 80% of OPCs are Ki67^+^ after 24 hours *in vitro* (B). Process length was determined using ImageJ. scOPC develop longer protrusions under proliferating conditions; unpaired t-test p<0.0001 (C). BrdU incorporation assay was performed and analyzed after different time points of fixation (6h, 30h or 54h); no significant difference in number of BrdU^+^ OPCs (D). *q*PCR did not show a significant difference in expression level of myelin- associated genes *Mbp, Plp1* and *Mag* (E-G). Exemplary brightfield images of differentiated cOL and scOL in 2D culture. Scale bar 100 µm (H). Quantification of morphology during differentiation is shown (I).

**Fig. 2-1** Validation of microarray candidate genes

Validation of differentially expressed genes where further verified by *q*PCR with original microarray probes (n = 3). Statistical analysis was performed with an unpaired Students’ t-test of each gene with significances indicated * p ≤ 0.05, ** p ≤ 0.01 and *** p ≤ 0.001 (A,B). New independent OPC samples isolated from P6-8 pups and cultured for a maximum of 6 days without passaging were analyzed by qPCR. *Skap2* (p = 0.0273), *Spata13* (p = 0.04), *Capn5* (p = 0.0166), but not *Nptx2* (p = 0.9515) were significantly upregulated in scOPC (C-F) while *Eya1* (p = 0.0122) and *Nr2e1* (p < 0.0001) but not, *Dbc-* 1 (p = 0.1367) and *Gria2* (p = 0.1234) were significantly downregulated in scOPC (G-J).

**Table 2-1** List of differentially expressed genes

List of differentially expressed gene probes sorted by probe rank and relative fold change using cOPC as reference group. Fold change indicates upregulation (>2) or downregulation (<0.5) of genes in scOPC compared to expression in cOPC. Differential expression was analyzed using direct hybridization analysis with quantile normalization, no background subtraction and the Illumina custom error model.

**Fig. 3-1** Differentiation of cOL and scOL after SKAP2 knockdown

Knockdown efficiency of *Skap2* during differentiation was evaluated in cOL (A) as well as scOL (B). *Mbp* and *Plp1* expression was not different at any time point in neither cOL (C) nor scOL (D) after *Skap2* knockdown. scOPC transduced with *Skap2* shRNA displayed a reduced number of sheath forming cells (53.82% ± 4.3 vs. 38.41% ± 3.1) and more cells remained mainly as multiple branched cells (36.39% ± 4.1 vs. 47.24% ± 2.9). Statictical analysis using Two-way ANOVA and Bonferroni post-test with * p < 0.05 (n = 3) (G).

**Fig. 3-2** Quantification of differentiation in tGFP^+^ cells after lentiviral transduction

Transduction efficiency was comparable for cOL^scr^ and cOL^Skap2sh^ determined by tGFP^+^ cells (68.85% ± 3.0 for cOL^scr^ and 73.12% ± 9.3 cOL^Skap2sh^) (A). Differentiation into O4^+^ cells was determined for tGFP^+^ cells in both conditions (58.79% ± 3.4 for cOL^scr^ and 53.8% ± 6.5 cOL^Skap2sh^) (B). Differentiation into MBP+ cells was not different in both conditions (17.39% ± 3.3 for cOL^scr^ and 13.7% ± 3.6 cOL^Skap2sh^) (C).

**Fig. 3-3** Lack of SKAP2 in scOPC affects myelin sheath formation but not myelin associated gene expression or proliferation

Myelin-associated genes *Mbp* and *Plp1* were not differentially expressed during differentiation in scOL^Skap2-/-^ compared to scOL^WT^ (A,B). Immunofluorescent staining with MBP, PDGFrα and DAPI was performed after 48 hours differentiation and no obvious difference in the number of MBP^+^ cells was determined (C,D). Number of Ki67^+^ cells was similar for scOPC^SKAP2-/-^ and scOPC^WT^ (E). The number of sheath forming cells was significantly reduced in scOL^Skap2-/-^ (18.22% ± 2.3) compared to scOL^WT^ (32.44% ± 3.8) (F). Statistical analysis was performed with Two-way ANOVA and Bonferroni post-test with * p < 0.05 (n = 3). Exemplary brightfield images display reduced numbers of scOL^Skap2-/-^ forming sheaths after 48 hours, scale bar 100 µm (G).

**Fig. 3-4** Lack of SKAP2 in cOL did not affect myelin-associated gene expression and proliferation

Myelin-associated genes *Mbp* and *Plp1* were not differentially expressed during differentiation in cOL^Skap2-/-^ compared to cOL^WT^ (A,B). Immunofluorescent staining with MBP, PDGFrα and DAPI was performed after 48 hours differentiation and no obvious difference in the number of MBP^+^ cells was determined (C,D). Number of Ki67^+^ cells was similar for cOPC^SKAP2-/-^ and cOPC^WT^ (E).

**Fig. 4-1** Knockdown efficiency of SKAP2 in cOPC and scOPC

Knockdown of *Skap2* in OPC reduced *Skap2* expression to 39.5% ± 7.1 in cOPC and 32.2% ± 1.8 in scOPC compared to scrambled control after 48 hours.

**Fig. 4-2** *SKAP2* expression in hiOL and knockdown efficiency of *hSKAP2* shRNA

SKAP2 expression increased significantly during differentiation from NPC to hiOL, unpaired t-test p = 0.0145 (A). Evaluation of *SKAP2* expression by *q*PCR showed a residual expression of 37.9% 72 hours after lentiviral knockdown (B)

## References

Alenghat FJ, Baca QJ, Rubin NT, Pao LI, Matozaki T, Lowell CA, Golan DE, Neel BG, Swanson KD (2012) Macrophages require Skap2 and Sirpalpha for integrin-stimulated cytoskeletal rearrangement. J Cell Sci 125: 5535–5545, doi:10.1242/jcs.111260.

Asazuma N, Wilde JI, Berlanga O, Leduc M, Leo A, Schweighoffer E, Tybulewicz V, Bon C, Liu SK, McGlade CJ, Schraven B, Watson SP (2000) Interaction of linker for activation of T cells with multiple adapter proteins in platelets activated by the glycoprotein VI-selective ligand, convulxin. J Biol Chem 275: 33427–33434, doi:10.1074/jbc.M001439200.

Ayoub E, Hall A, Scott AM, Chagnon MJ, Miquel G, Halle M, Noda M, Bikfalvi A, Tremblay ML (2013) Regulation of the Src Kinase-associated Phosphoprotein 55 Homologue by the Protein Tyrosine Phosphatase PTP-PEST in the Control of Cell Motility. J Biol Chem 288: 25739– 25748, doi:10.1074/jbc.M113.501007.

Barateiro A, Fernandes A (2014) Temporal oligodendrocyte lineage progression: In vitro models of proliferation, differentiation and myelination. Biochim Biophys Acta - Mol Cell Res 1843: 1917–1929, doi:https://doi.org/10.1016/j.bbamcr.2014.04.018.

Bauer NG, Richter-Landsberg C, Ffrench-Constant C (2009) Role of the Oligodendroglial Cytoskeleton in Differentiation and Myelination. Glia 57: 1691–1705, doi:10.1002/glia.20885.

Bechler ME, Byrne L, Ffrench-Constant C (2015) CNS Myelin Sheath Lengths Are an Intrinsic Property of Oligodendrocytes. Curr Biol 25: 2411–2416, doi:10.1016/j.cub.2015.07.056.

Boras M, Volmering S, Bokemeyer A, Rossaint J, Block H, Bardel B, Van Marck V, Heitplatz B, Kliche S, Reinhold A, Lowell C, Zarbock A (2017) Skap2 is required for beta(2) integrin-mediated neutrophil recruitment and functions. J Exp Med 214: 851–874, doi:10.1084/jem.20160647.

Bourette RP, Thérier J, Mouchiroud G (2005) Macrophage colony-stimulating factor receptor induces tyrosine phosphorylation of SKAP55R adaptor and its association with actin. Cell Signal 17: 941–949, doi:10.1016/j.cellsig.2004.11.009.

Brown TL, Macklin WB (2020) The Actin Cytoskeleton in Myelinating Cells. Neurochem Res 45: 684–693, doi:10.1007/s11064-019-02753-0.

van Bruggen D, Agirre E, Castelo-Branco G (2017) Single-cell transcriptomic analysis of oligodendrocyte lineage cells. Curr Opin Neurobiol 47: 168–175, doi:10.1016/j.conb.2017.10.005.

Bureau J-F, Cassonnet P, Grange L, Dessapt J, Jones L, Demeret C, Sakuntabhai A, Jacob Y (2018) The SRC-family tyrosine kinase HCK shapes the landscape of SKAP2 interactome. Oncotarget 9: 13102–13115, doi:10.18632/oncotarget.24424.

de Castro F, Bribián A, Ortega MC (2013) Regulation of oligodendrocyte precursor migration during development, in adulthood and in pathology. Cell Mol Life Sci 70: 4355–4368, doi:10.1007/s00018-013-1365-6.

Ehrlich M, Mozafari S, Glatza M, Starost L, Velychko S, Hallmann A-L, Cui Q-L, Schambach A, Kim K-P, Bachelin C, Marteyn A, Hargus G, Johnson RM, Antel J, Sterneckert J, Zaehres H, Schöler HR, Baron-Van Evercooren A, Kuhlmann T (2017) Rapid and efficient generation of oligodendrocytes from human induced pluripotent stem cells using transcription factors. Proc Natl Acad Sci doi:10.1073/pnas.1614412114.

Feng J-F, Gao X-F, Pu Y, Burnstock G, Xiang Z, He C (2015) P2X7 receptors and Fyn kinase mediate ATP-induced oligodendrocyte progenitor cell migration. Purinergic Signal 11: 361– 369, doi:10.1007/s11302-015-9458-3.

Franklin RJM, Goldman SA (2015) Glia Disease and Repair-Remyelination. Cold Spring Harb Perspect Biol 7: a020594, doi:10.1101/cshperspect.a020594.

Gailit J, Ruoslahti E (1988) Regulation of the fibronectin receptor affinity by divalent cations. J Biol Chem 263: 12927–12932, doi:10.1016/S0021-9258(18)37650-6.

Hill RA, Patel KD, Medved J, Reiss AM, Nishiyama A (2013) NG2 cells in white matter but not gray matter proliferate in response to PDGF. J Neurosci 33: 14558–14566, doi:10.1523/JNEUROSCI.2001-12.2013.

Horiuchi M, Suzuki-Horiuchi Y, Akiyama T, Itoh A, Pleasure D, Carstens E, Itoh T (2017) Differing intrinsic biological properties between forebrain and spinal oligodendroglial lineage cells. J Neurochem 142: 378–391, doi:10.1111/jnc.14074.

Katanov C, Novak N, Vainshtein A, Golani O, Dupree JL, Peles E (2020) N-Wasp Regulates Oligodendrocyte Myelination. J Neurosci 40: 6103 LP – 6111, doi:10.1523/JNEUROSCI.0912-20.2020.

Kessaris N, Fogarty M, Iannarelli P, Grist M, Wegner M, Richardson WD (2006) Competing waves of oligodendrocytes in the forebrain and postnatal elimination of an embryonic lineage. Nat Neurosci 9: 173–179, doi:10.1038/nn1620.

Klein C, Kramer EM, Cardine AM, Schraven B, Brandt R, Trotter J (2002) Process outgrowth of oligodendrocytes is promoted by interaction of Fyn kinase with the cytoskeletal protein Tau. J Neurosci 22: 698–707, doi:10.1523/jneurosci.22-03-00698.2002.

Königsberger S, Peckl-Schmid D, Zaborsky N, Patzak I, Kiefer F, Achatz G (2010) HPK1 associates with SKAP-HOM to negatively regulate Rap1-mediated B-lymphocyte adhesion. PLoS One 5: doi:10.1371/journal.pone.0012468.

Kuhn S, Gritti L, Crooks D, Dombrowski Y (2019) Oligodendrocytes in Development, Myelin Generation and Beyond. Cells 8: doi:10.3390/cells8111424.

Lentferink DH, Jongsma JM, Werkman I, Baron W (2018) Grey matter OPCs are less mature and less sensitive to IFN gamma than white matter OPCs: consequences for remyelination. Sci Rep 8: 2113, doi:10.1038/s41598-018-19934-6.

Livak KJ, Schmittgen TD (2001) Analysis of relative gene expression data using real-time quantitative PCR and the 2(T)(-Delta Delta C) method. Methods 25: 402–408, doi:10.1006/meth.2001.1262.

Marie-Cardine A, Verhagen AM, Eckerskorn C, Schraven B (1998) SKAP-HOM, a novel adaptor protein homologous to the FYN-associated protein SKAP55. FEBS Lett 435: 55–60, doi:10.1016/S0014-5793(98)01040-0.

Marques S, van Bruggen D, Vanichkina DP, Floriddia EM, Munguba H, Väremo L, Giacomello S, Falcão AM, Meijer M, Björklund ÅK, Hjerling-Leffler J, Taft RJ, Castelo-Branco G (2018) Transcriptional Convergence of Oligodendrocyte Lineage Progenitors during Development. Dev Cell 46: 504–517.e7, doi:10.1016/j.devcel.2018.07.005.

Marques S, Zeisel A, Codeluppi S, van Bruggen D, Falcao AM, Xiao L, Li H, Haring M, Hochgerner H, Romanov RA, Gyllborg D, Munoz-Manchado AB, La Manno G, Lonnerberg P, Floriddia EM, Rezayee F, Ernfors P, Arenas E, Hjerling-Leffler J, Harkany T, Richardson WD, Linnarsson S, Castelo-Branco G (2016) Oligodendrocyte heterogeneity in the mouse juvenile and adult central nervous system. Science (80-) 352: 1326–1329, doi:10.1126/science.aaf6463.

Meijering E, Dzyubachyk O, Smal I (2012) Methods for cell and particle tracking. Methods Enzymol 504: 183–200, doi:10.1016/B978-0-12-391857-4.00009-4.

Miguez A, Ducret S, Di Meglio T, Parras C, Hmidan H, Haton C, Sekizar S, Mannioui A, Vidal M, Kerever A, Nyabi O, Haigh J, Zalc B, Rijli FM, Thomas JL (2012) Opposing roles for Hoxa2 and Hoxb2 in hindbrain oligodendrocyte patterning. J Neurosci 32: 17172–17185, doi:10.1523/JNEUROSCI.0885-12.2012.

Miyamoto Y, Yamauchi J, Tanoue A (2008) Cdk5 Phosphorylation of WAVE2 Regulates Oligodendrocyte Precursor Cell Migration through Nonreceptor Tyrosine Kinase Fyn. J Neurosci 28: 8326–8337, doi:10.1523/JNEUROSCI.1482-08.2008.

Morita S, Kojima T, Kitamura T (2000) Plat-E: an efficient and stable system for transient packaging of retroviruses. Gene Ther 7: 1063–1066, doi:10.1038/sj.gt.3301206.

Musah AS, Brown TL, Jeffries MA, Shang Q, Hashimoto H, Evangelou A V, Kowalski A, Batish M, Macklin WB, Wood TL (2020) Mechanistic Target of Rapamycin Regulates the Oligodendrocyte Cytoskeleton during Myelination. J Neurosci 40: 2993–3007, doi:10.1523/JNEUROSCI.1434-18.2020.

Nawaz S, Sanchez P, Schmitt S, Snaidero N, Mitkovski M, Velte C, Bruckner BR, Alexopoulos I, Czopka T, Jung SY, Rhee JS, Janshoff A, Witke W, Schaap IAT, Lyons DA, Simons M (2015) Actin filament turnover drives leading edge growth during myelin sheath formation in the central nervous system. Dev Cell 34: 139–151, doi:10.1016/j.devcel.2015.05.013.

Nieberler M, Reuning U, Reichart F, Notni J, Wester H-J, Schwaiger M, Weinmüller M, Räder A, Steiger K, Kessler H (2017) Exploring the Role of RGD-Recognizing Integrins in Cancer. Cancers (Basel) 9: 116, doi:10.3390/cancers9090116.

O’Meara RW, Michalski JP, Kothary R (2011) Integrin Signaling in Oligodendrocytes and Its Importance in CNS Myelination. J Signal Transduct 2011: 10.1155/2011/354091, doi:10.1155/2011/354091.

Pawlowski M, Ortmann D, Bertero A, Tavares JM, Pedersen RA, Vallier L, Kotter MRN (2017) Inducible and Deterministic Forward Programming of Human Pluripotent Stem Cells into Neurons, Skeletal Myocytes, and Oligodendrocytes. Stem cell reports 8: 803–812, doi:10.1016/j.stemcr.2017.02.016.

Philippidou P, Dasen JS (2013) Hox Genes: Choreographers in Neural Development, Architects of Circuit Organization. Neuron 80: 12–34, doi:10.1016/j.neuron.2013.09.020.

Pierschbacher MD, Ruoslahti E (1984) Cell attachment activity of fibronectin can be duplicated by small synthetic fragments of the molecule. Nature 309: 30–33, doi:10.1038/309030a0.

Reinhardt P, Glatza M, Hemmer K, Tsytsyura Y, Thiel CS, Höing S, Moritz S, Parga JA, Wagner L, Bruder JM, Wu G, Schmid B, Röpke A, Klingauf J, Schwamborn JC, Gasser T, Schöler HR, Sterneckert J (2013) Derivation and expansion using only small molecules of human neural progenitors for neurodegenerative disease modeling. PLoS One 8: e59252– e59252, doi:10.1371/journal.pone.0059252.

Schambach A, Galla M, Modlich U, Will E, Chandra S, Reeves L, Colbert M, Williams DA, von Kalle C, Baum C (2006) Lentiviral vectors pseudotyped with murine ecotropic envelope: increased biosafety and convenience in preclinical research. Exp Hematol 34: 588–592, doi:10.1016/j.exphem.2006.02.005.

Schilling TF, Knight RD (2001) Origins of anteroposterior patterning and Hox gene regulation during chordate evolution. Philos Trans R Soc LondonSeries B 356: 1599–1613, doi:10.1098/rstb.2001.0918.

Shek DT, Ma CM (2011) Longitudinal data analyses using linear mixed models in SPSS: concepts, procedures and illustrations. ScientificWorldJournal 11: 42–76, doi:10.1100/tsw.2011.2.

Shimamura S, Sasaki K, Tanaka M (2013) The Src Substrate SKAP2 Regulates Actin Assembly by Interacting with WAVE2 and Cortactin Proteins. J Biol Chem 288: 1171–1183, doi:10.1074/jbc.M112.386722.

Simons M, Nave K-A (2015) Oligodendrocytes: Myelination and Axonal Support. Cold Spring Harb Perspect Biol 8: a020479, doi:10.1101/cshperspect.a020479.

Smith JR, Maguire S, Davis LA, Alexander M, Yang F, Chandran S, ffrench-Constant C, Pedersen RA (2008) Robust, persistent transgene expression in human embryonic stem cells is achieved with AAVS1-targeted integration. Stem Cells 26: 496–504, doi:10.1634/stemcells.2007-0039.

Snaidero N, Velte C, Myllykoski M, Raasakka A, Ignatev A, Werner HB, Erwig MS, Mobius W, Kursula P, Nave KA, Simons M (2017) Antagonistic Functions of MBP and CNP Establish Cytosolic Channels in CNS Myelin. Cell Rep 18: 314–323, doi:S2211-1247(16)31760-0.

Sperber BR, McMorris FA (2001) Fyn tyrosine kinase regulates oligodendroglial cell development but is not required for morphological differentiation of oligodendrocytes. J Neurosci Res 63: 303–312, doi:10.1002/1097-4547(20010215)63:4&303::AID-JNR1024>3.0.CO;2-A.

Takuma K, Carina H (2017) Evolutionary conservation and conversion of Foxg1 function in brain development. Dev Growth Differ 59: 258–269, doi:10.1111/dgd.12367.

Tanaka M, Shimamura S, Kuriyama S, Maeda D, Goto A, Aiba N (2016) SKAP2 Promotes Podosome Formation to Facilitate Tumor-Associated Macrophage Infiltration and Metastatic Progression. Cancer Res 76: 358–369, doi:10.1158/0008-5472.CAN-15-1879.

Thomason EJ, Escalante M, Osterhout DJ, Fuss B (2020) The oligodendrocyte growth cone and its actin cytoskeleton: A fundamental element for progenitor cell migration and CNS myelination. Glia 68: 1329–1346, doi:10.1002/glia.23735.

Togni M, Swanson KD, Reimann S, Kliche S, Pearce AC, Simeoni L, Reinhold D, Wienands J, Neel BG, Schraven B, Gerber A (2005) Regulation of in vitro and in vivo immune functions by the cytosolic adaptor protein SKAP-HOM. Mol Cell Biol 25: 8052–8063, doi:10.1128/MCB.25.18.8052-8063.2005.

Tsai HH, Niu J, Munji R, Davalos D, Chang J, Zhang H, Tien AC, Kuo CJ, Chan JR, Daneman R, Fancy SP (2016) Oligodendrocyte precursors migrate along vasculature in the developing nervous system. Science 351: 379–384, doi:10.1126/science.aad3839.

Tsuneoka Y, Tsukahara S, Yoshida S, Takase K, Oda S, Kuroda M, Funato H (2017) Moxd1 Is a Marker for Sexual Dimorphism in the Medial Preoptic Area, Bed Nucleus of the Stria Terminalis and Medial Amygdala. Front Neuroanat 11: 26, doi:10.3389/fnana.2017.00026.

Tumpel S, Wiedemann LM, Krumlauf R (2009) Hox genes and segmentation of the vertebrate hindbrain. Curr Top Dev Biol 88: 103–137, doi:10.1016/S0070-2153(09)88004-6.

Tyler WA, Jain MR, Cifelli SE, Li Q, Ku L, Feng Y, Li H, Wood TL (2011) Proteomic identification of novel targets regulated by the mammalian target of rapamycin pathway during oligodendrocyte differentiation. Glia 59: 1754–1769, doi:10.1002/glia.21221.

Vigano F, Mobius W, Gotz M, Dimou L (2013) Transplantation reveals regional differences in oligodendrocyte differentiation in the adult brain. Nat Neurosci 16: 1370–1372, doi:10.1038/nn.3503.

Watkins TA, Emery B, Mulinyawe S, Barres BA (2008) Distinct Stages of Myelination Regulated by gamma-Secretase and Astrocytes in a Rapidly Myelinating CNS Coculture System. Neuron 60: 555–569, doi:10.1016/j.neuron.2008.09.011.

Xu YKT, Chitsaz D, Brown RA, Cui QL, Dabarno MA, Antel JP, Kennedy TE (2019) Deep learning for high-throughput quantification of oligodendrocyte ensheathment at single-cell resolution. Commun Biol 2: 116, doi:10.1038/s42003-019-0356-z.

Zhou L, Zhang Z, Zheng Y, Zhu Y, Wei Z, Xu H, Tang Q, Kong X, Hu L (2011) SKAP2, a novel target of HSF4b, associates with NCK2/F-actin at membrane ruffles and regulates actin reorganization in lens cell. J Cell Mol Med 15: 783–795, doi:10.1111/j.1582-4934.2010.01048.x.

Zuchero JB, Fu M-M, Sloan SA, Ibrahim A, Olson A, Zaremba A, Dugas JC, Wienbar S, Caprariello A V, Kantor C, Leonoudakis D, Lariosa-Willingham K, Kronenberg G, Gertz K, Soderling SH, Miller RH, Barres BA (2015) CNS myelin wrapping is driven by actin disassembly. Dev Cell 34: 152–167, doi:10.1016/j.devcel.2015.06.011.

