## Supplementary figures and images for "SKAP2 as a new regulator of oligodendroglial migration and myelin sheath formation"

### Figure 1-1

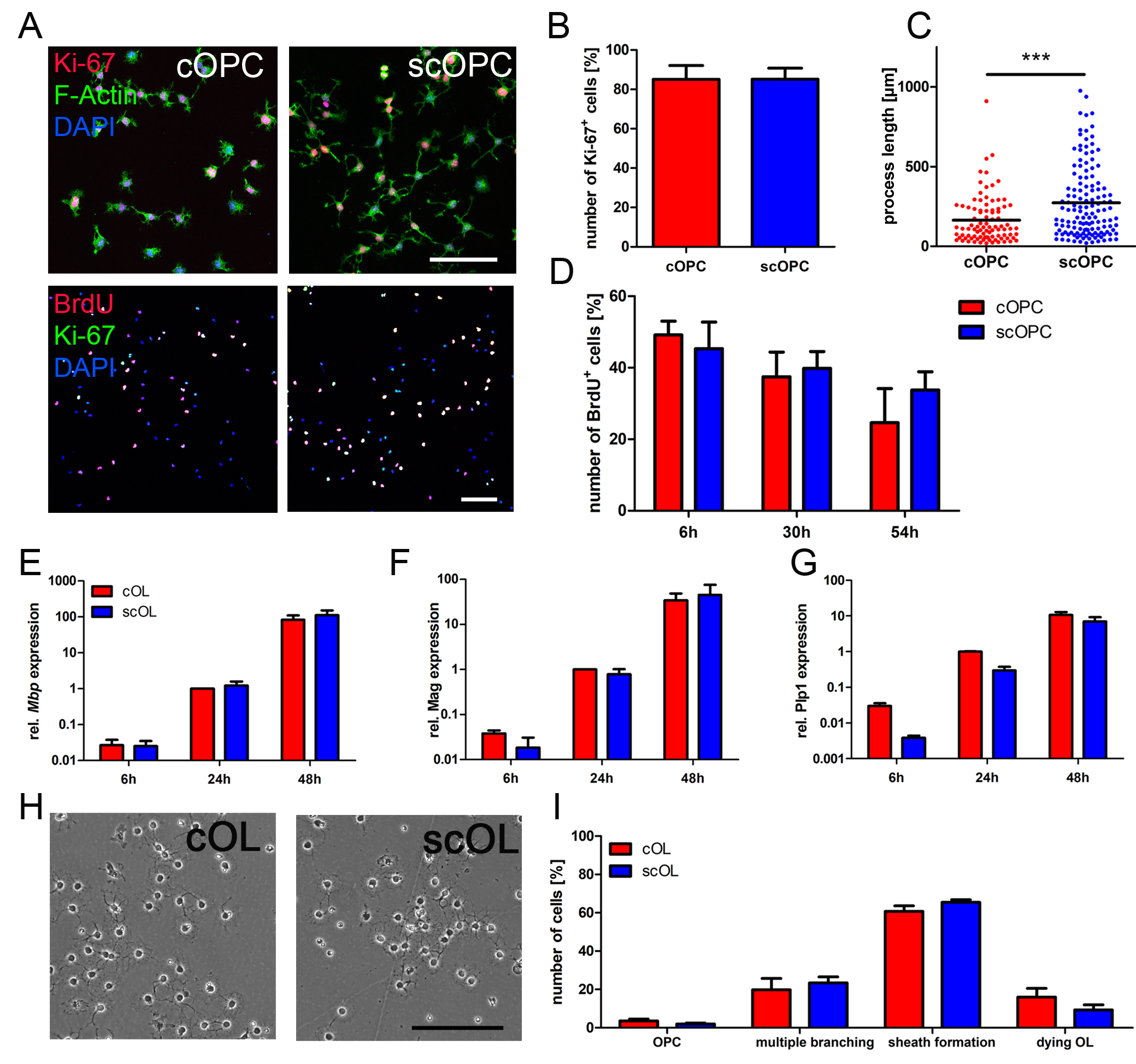

### Figure 2-1

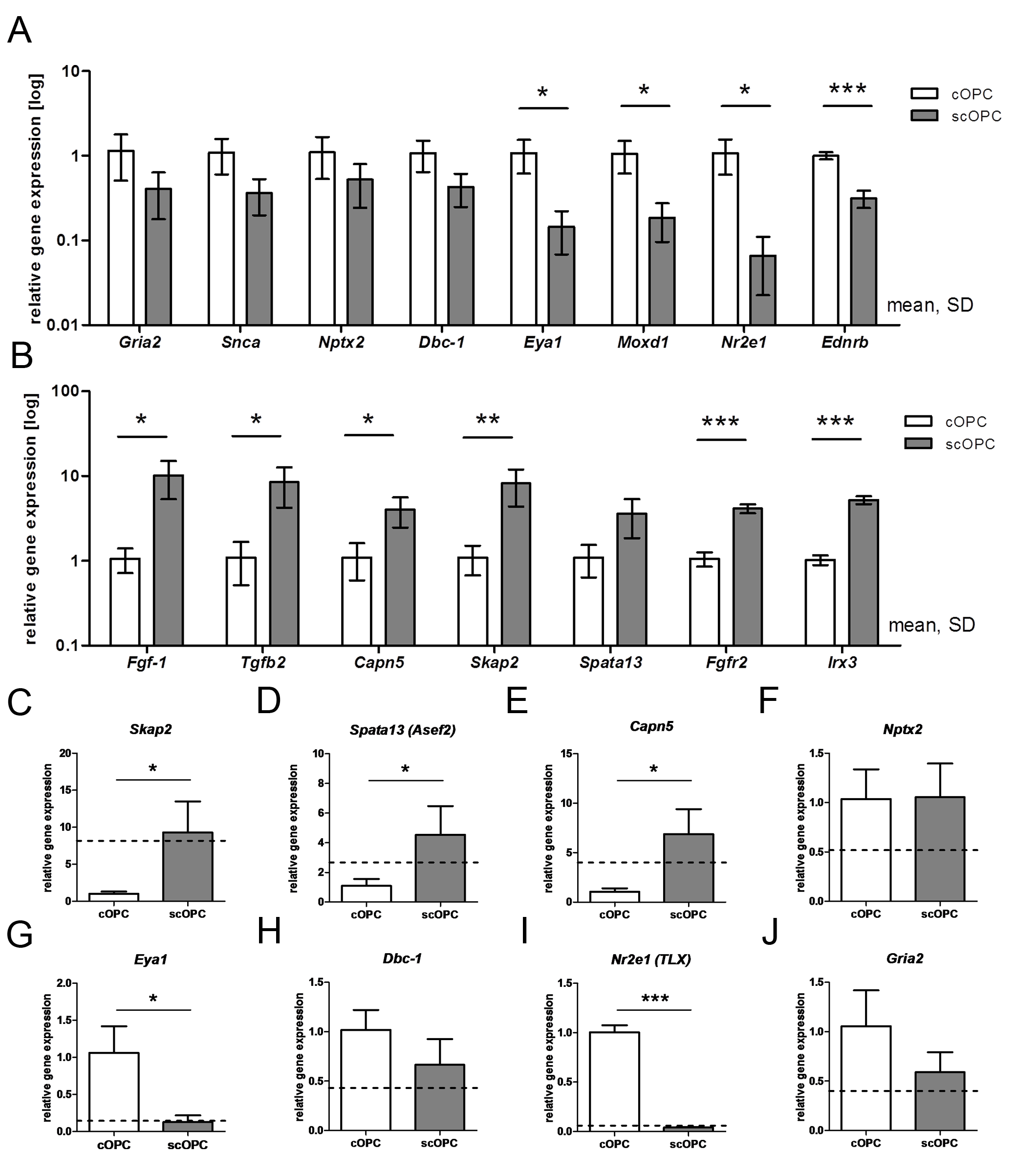

### Figure 3-1

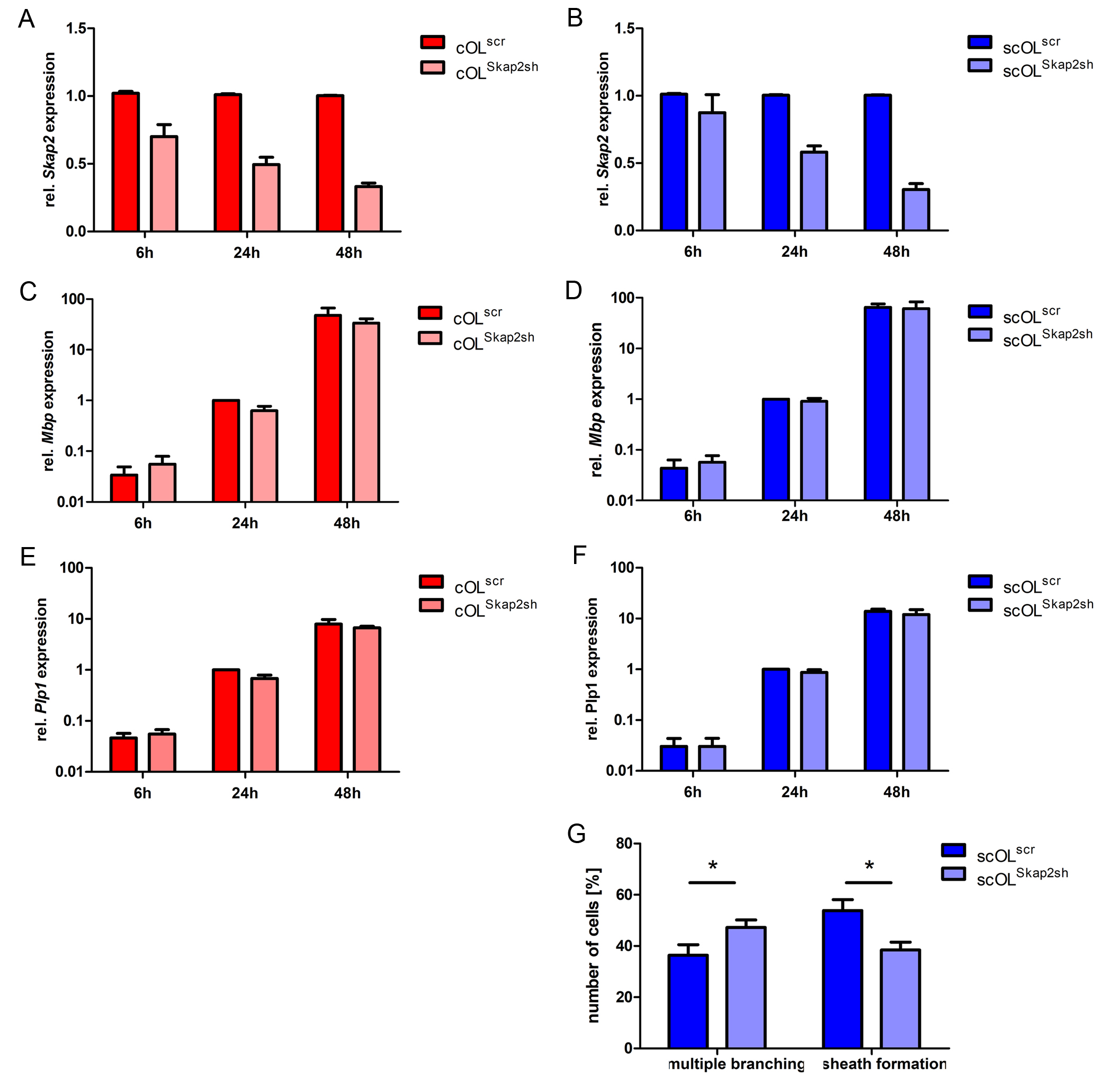

### Figure 3-2

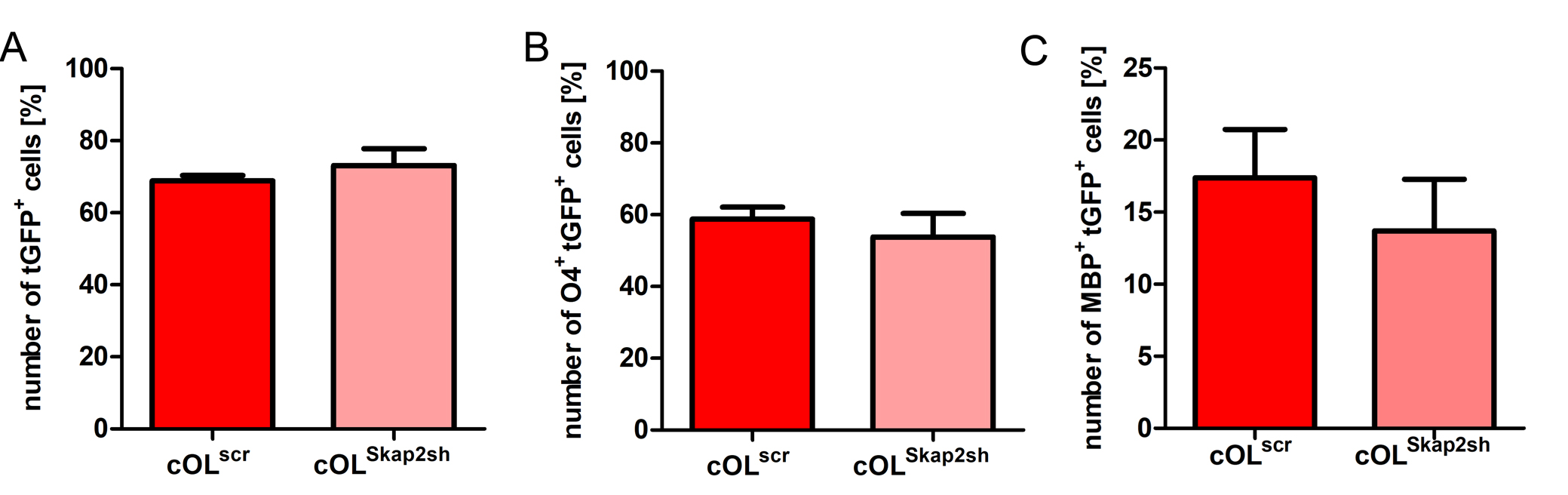

### Figure 3-3

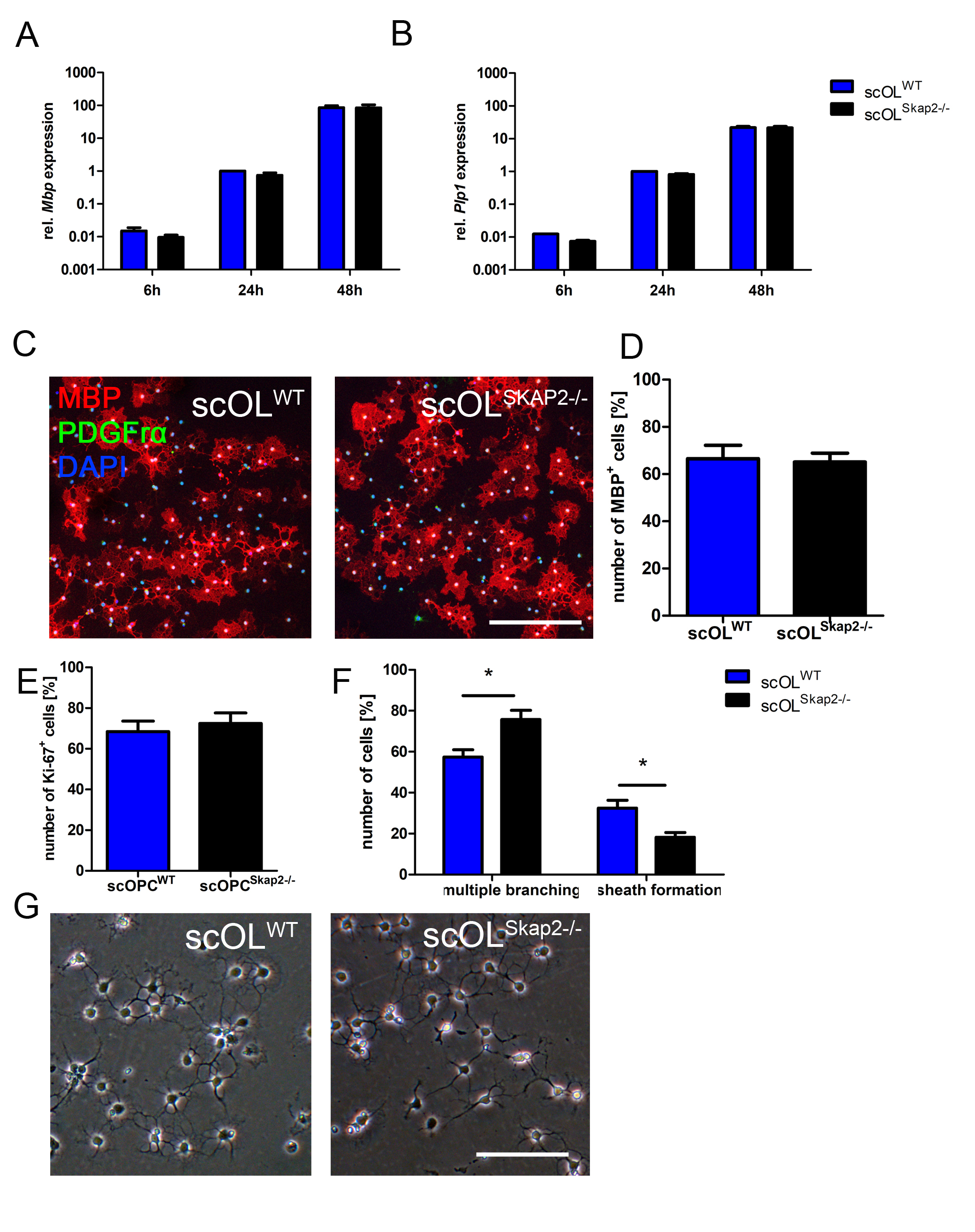

### Figure 3-4

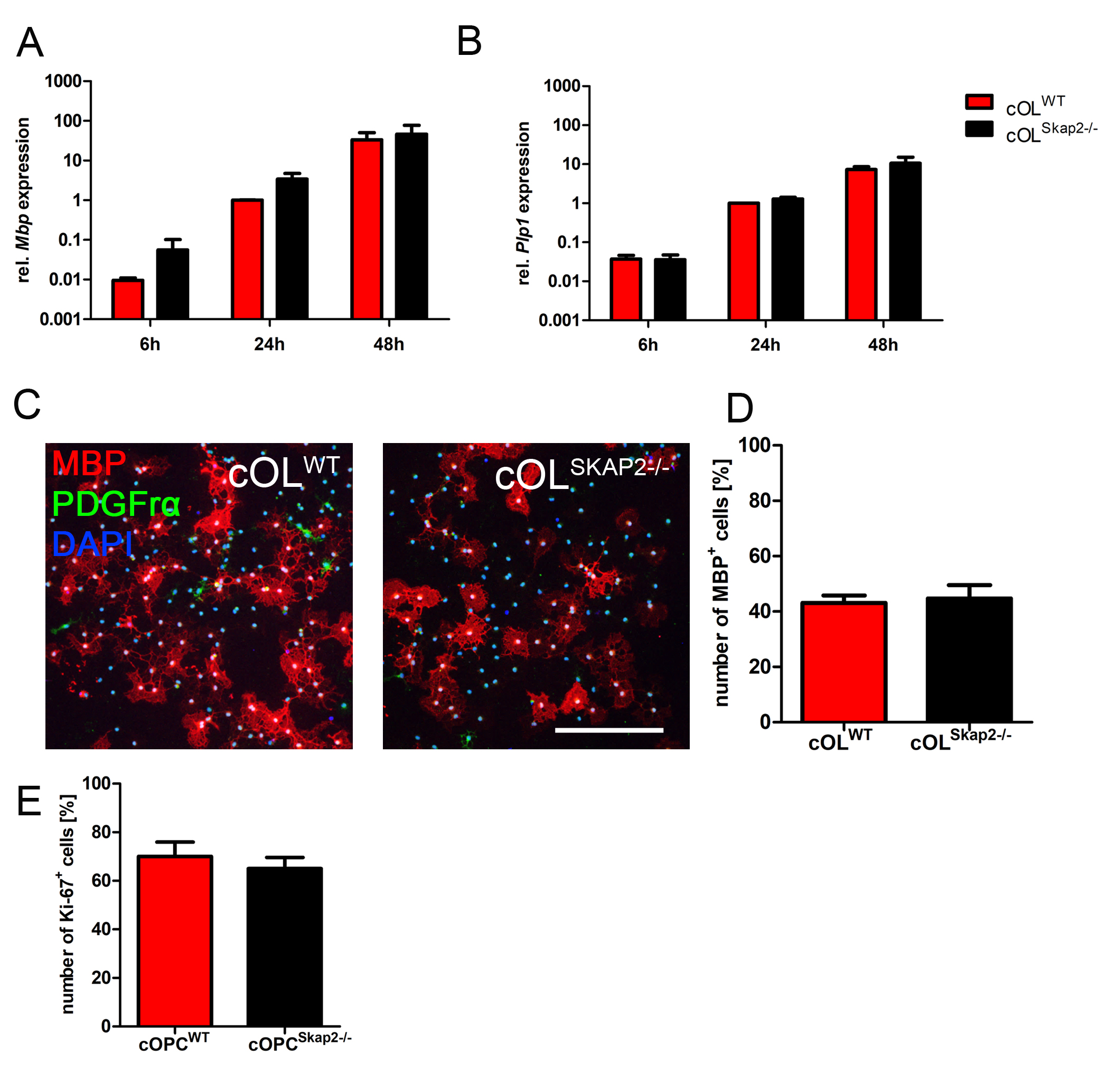

### Figure 4-1

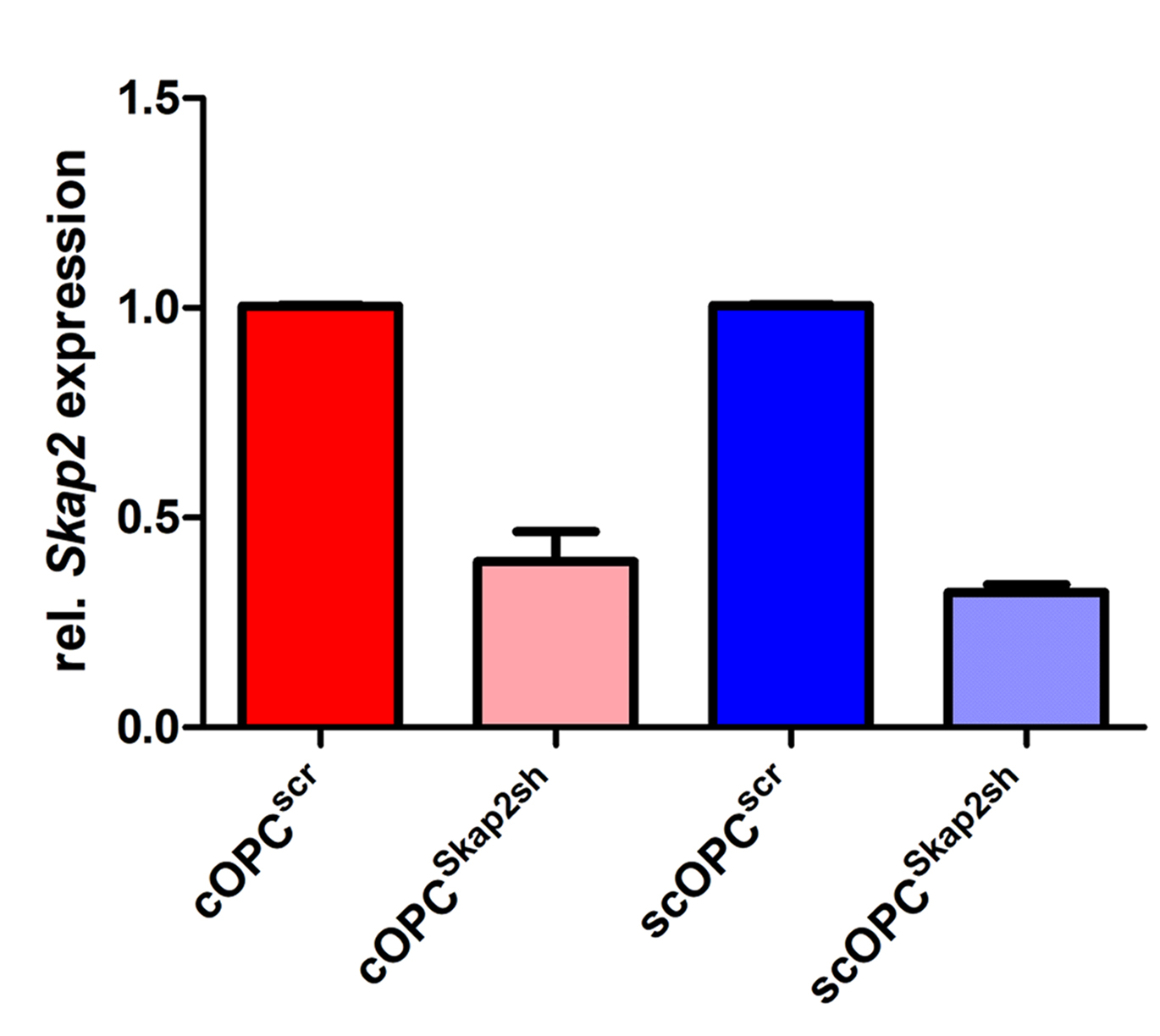

### Figure 4-2

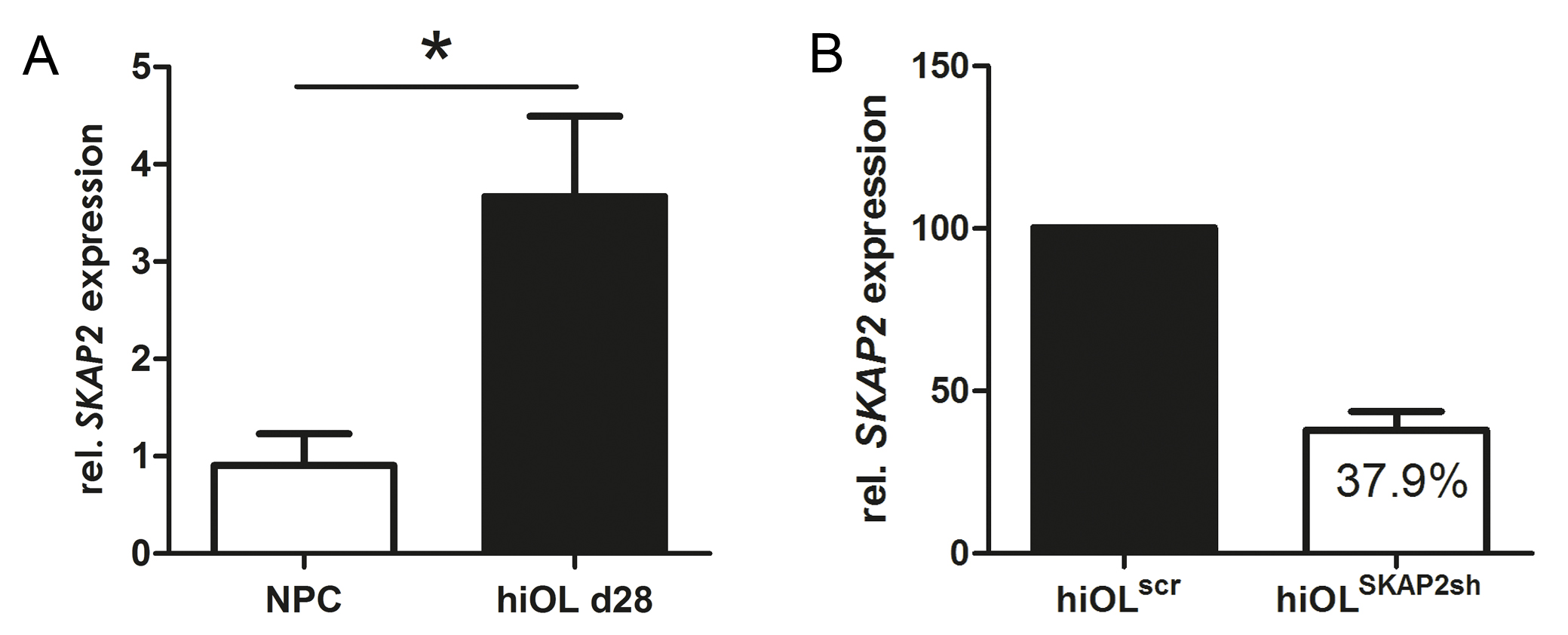
