## Supplementary material for "SKAP2 as a new regulator of oligodendroglial migration and myelin sheath formation": Table 2-1

| rank | probe | fold change | rank | probe | fold change |
| --- | --- | --- | --- | --- | --- |
| 1 | *Hoxb2* | 6.860 | 1 | *Moxd1* | 0.176 |
| 2 | *Adcy8* | 5.662 | 2 | *Moxd1* | 0.187 |
| 3 | *Irx5* | 5.156 | 3 | *Eya1* | 0.204 |
| 4 | *Tmem132c* | 4.720 | 4 | *Gpr149* | 0.229 |
| 5 | *Irx3* | 4.484 | 5 | *Chst7* | 0.251 |
| 6 | *Fgf1* | 4.301 | 6 | *Si* | 0.294 |
| 7 | *5830411I20* | 3.985 | 7 | *Atp10b* | 0.297 |
| 8 | *Btbd14a* | 3.890 | 8 | *Liph* | 0.333 |
| 9 | *Skap2* | 3.573 | 9 | *Snca* | 0.338 |
| 10 | *Fgfr2* | 3.553 | 10 | *Liph* | 0.342 |
| 11 | *Hoxc10* | 3.541 | 11 | *Snca* | 0.346 |
| 12 | *Capn5* | 3.363 | 12 | *Sox9* | 0.382 |
| 13 | *6720458D17Rik* | 3.104 | 13 | *A730017C20Rik* | 0.388 |
| 14 | *Akap7* | 3.067 | 14 | *Egr3* | 0.410 |
| 15 | *Tgfb2* | 3.053 | 15 | *Dbc1* | 0.413 |
| 16 | *Pdzrn3* | 2.997 | 16 | *Gria2* | 0.416 |
| 17 | *Efna5* | 2.899 | 17 | *Ednrb* | 0.423 |
| 18 | *Gm687* | 2.818 | 18 | *Ednrb* | 0.435 |
| 19 | *Hoxa4* | 2.705 | 19 | *Plxnb3* | 0.436 |
| 20 | *9630031F12Rik* | 2.704 | 20 | *Mest* | 0.437 |
| 21 | *LOC100047808* | 2.619 | 21 | *Nr2e1* | 0.439 |
| 22 | *Irx2* | 2.461 | 22 | *Ckmt1* | 0.440 |
| 23 | *Msx1* | 2.458 | 23 | *Dbc1* | 0.450 |
| 24 | *Sdccag33l* | 2.444 | 24 | *Cacna2d1* | 0.461 |
| 25 | *Irx2* | 2.403 | 25 | *Luzp2* | 0.466 |
| 26 | *Capn6* | 2.386 | 26 | *Nptx2* | 0.474 |
| 27 | *4933436C20Rik* | 2.360 | 27 | *Itpkb* | 0.479 |
| 28 | *Whrn* | 2.350 | 28 | *Gria2* | 0.482 |
| 29 | *Rab26* | 2.301 | 29 | *Mest* | 0.484 |
| 30 | *D12Ertd553e* | 2.277 | 30 | *Lypd6* | 0.487 |
| 31 | *Frmd4a* | 2.275 | 31 | *2610017I09Rik* | 0.489 |
| 32 | *Cdkn1c* | 2.270 | 32 | *Igfbp2* | 0.490 |
| 33 | *2610307O08Rik* | 2.268 | 33 | *Lypd6* | 0.490 |
| 34 | *Cpne8* | 2.268 | 34 | *Igfbp2* | 0.494 |
| 35 | *Ndrg2* | 2.254 | 35 | *4833424O15Rik* | 0.495 |
| 36 | *Rit2* | 2.254 |  |  |  |
| 37 | *Tshz2* | 2.217 |  |  |  |
| 38 | *Mbp* | 2.210 |  |  |  |
| 39 | *Spata13* | 2.205 |  |  |  |
| 40 | *Tsc22d1* | 2.198 |  |  |  |
| 41 | *D230005E09Rik* | 2.177 |  |  |  |
| 42 | *Lgr5* | 2.177 |  |  |  |
| 43 | *Cpne8* | 2.173 |  |  |  |
| 44 | *Eln* | 2.142 |  |  |  |
| 45 | *Arpp21* | 2.137 |  |  |  |
| 46 | *Grb10* | 2.131 |  |  |  |
| 47 | *Lgals1* | 2.131 |  |  |  |
| 48 | *Rassf3* | 2.125 |  |  |  |
| 49 | *St8sia1* | 2.085 |  |  |  |
| 50 | *Timp2* | 2.020 |  |  |  |
| 51 | *Anxa2* | 2.015 |  |  |  |
|  | **Upregulation** | **50 genes** |  | **Downregulation** | **28 genes** |
